# Axon guidance during CNS regeneration is required for specific brain innervation

**DOI:** 10.1101/2023.09.04.556244

**Authors:** Céline Delpech, Julia Schaeffer, Noemie Vilallongue, Amin Benadjal, Beatrice Blot, Blandine Excoffier, Elise Plissonnier, Floriane Albert, Antoine Paccard, Yvrick Zagar, Valérie Castellani, Stephane Belin, Alain Chédotal, Homaira Nawabi

**Author notes:** These authors contributed equally to the work and are listed in alphabetical order (Céline Delpech, Julia Schaeffer, Noemie Vilallongue).

## Abstract

Reconstruction of functional neuronal circuits in the mature brain remains a big challenge in the field of central nervous system (CNS) repair. Despite achievement of robust, long-distance regeneration through modulation of specific neuronal intrinsic growth properties, functional recovery is still limited due to major guidance defects of regenerating axons. Using co-activation of mTOR, JAK/STAT and c-myc pathways in retinal ganglion cells (RGC), we highlight that regenerating axons avoid the suprachiasmatic nucleus (SCN) due to repulsive mechanisms. We show that Slit/Robo guidance signaling is responsible for this reinnervation failure. In vivo suppression of this repulsive signaling allows regenerating axons to enter the SCN. The newly formed circuit is associated with functional behavioral recovery. Our results provide evidence that axon guidance mechanisms are required in the context of mature neuronal circuit repair.

## Main text

In adult mammals, central nervous system (CNS) neurons are unable to regenerate, leading to permanent disabilities in patients with CNS injury. Using the mouse visual system as a gold-standard experimental model, axon regeneration of retinal ganglion cells (RGC) was unlocked through the modulation of developmentally- or injury-regulated pathways, such as mTOR, and JAK/STAT and c-myc ^1–3^. Moreover, the co-modulation of such pathways leads to long-distance regeneration, where RGC axons can regrow from the eye to the brain ^3–6^. However, no functional recovery has been achieved as few regenerating RGC axons reach their proper targets due to major guidance defect. Interestingly, this has been observed across all models of regeneration ^7–10^. Therefore, not only do axons fail to reconnect with their proper target, but aberrant axonal projections can form deleterious circuits that potentially impede functional recovery ^11^. Thus, a new challenge stands out: how to drive regenerating axons to their correct target(s) in order to restore function?

Axon guidance is a process well described during development, when billions of neurons extend axons that reach and make connections with target cells throughout the body ^12^. This process is governed by environmental guidance cues, which shape neuronal circuits by interacting with receptors expressed on the growth cone of developing axons. Yet, a growing body of evidence strongly suggests that guidance mechanisms also play a key role in adult CNS regeneration ^13,14^. Our recent work shows that many guidance cues are expressed in the mature visual system, and that most of them have a repulsive activity during development. Surprisingly, regenerating axons are able to respond to these cues, thereby opening the possibility to control their trajectories ^15^. In this study, we demonstrate that brain retinorecipient nuclei express repulsive cues that impair reinnervation after injury. As a proof-of-concept, we focus on the functional reinnervation of the suprachiasmatic nucleus (SCN), the master regulator of circadian rhythms ^16^. Our results show that Slit/Robo signaling controls SCN reinnervation by regenerating axons.

To induce a long-distance regeneration of RGC axons, we co-activated mTOR, JAK/STAT and c-myc pathways, via injection of AAV2-Cre/CNTF/c-myc in one eye of Pten^fl/fl^ SOCS3^fl/fl^ mice, followed by optic nerve crush injury (ONC) (**Figure 1A-B**) ^3^. 28 days post-crush (28dpc), RGC axons display robust regeneration along the optic nerve, up to the optic chiasm and beyond (**Figure 1B**), as described before ^3^. In intact wild-type condition (pink-labeled tract, **Figure 1C**), 95% of RGC axons cross the midline to project contralaterally, while 5% project into the ipsilateral tract (**Figure 1D**) ^17^. However, during regeneration, we observed multiple guidance defects at the optic chiasm and in the SCN region (**Figure 1C**). For example, 74% axons do not cross the midline, whereas 26% project contralaterally. In addition, 10% grow into the contralateral optic nerve (**Figure 1E**). Strikingly, when we looked at the SCN region, we found that regenerating axons grow around and in between the bilaterally symmetrical nuclei, but fail to enter them (**Figure 1F**). In intact condition, the SCN receives equal innervation from both eyes (pink-labeled tract, **Figure 1C, F**). Therefore, we asked whether the intact contralateral RGC axons could interfere with SCN reinnervation.

**Figure 1:**
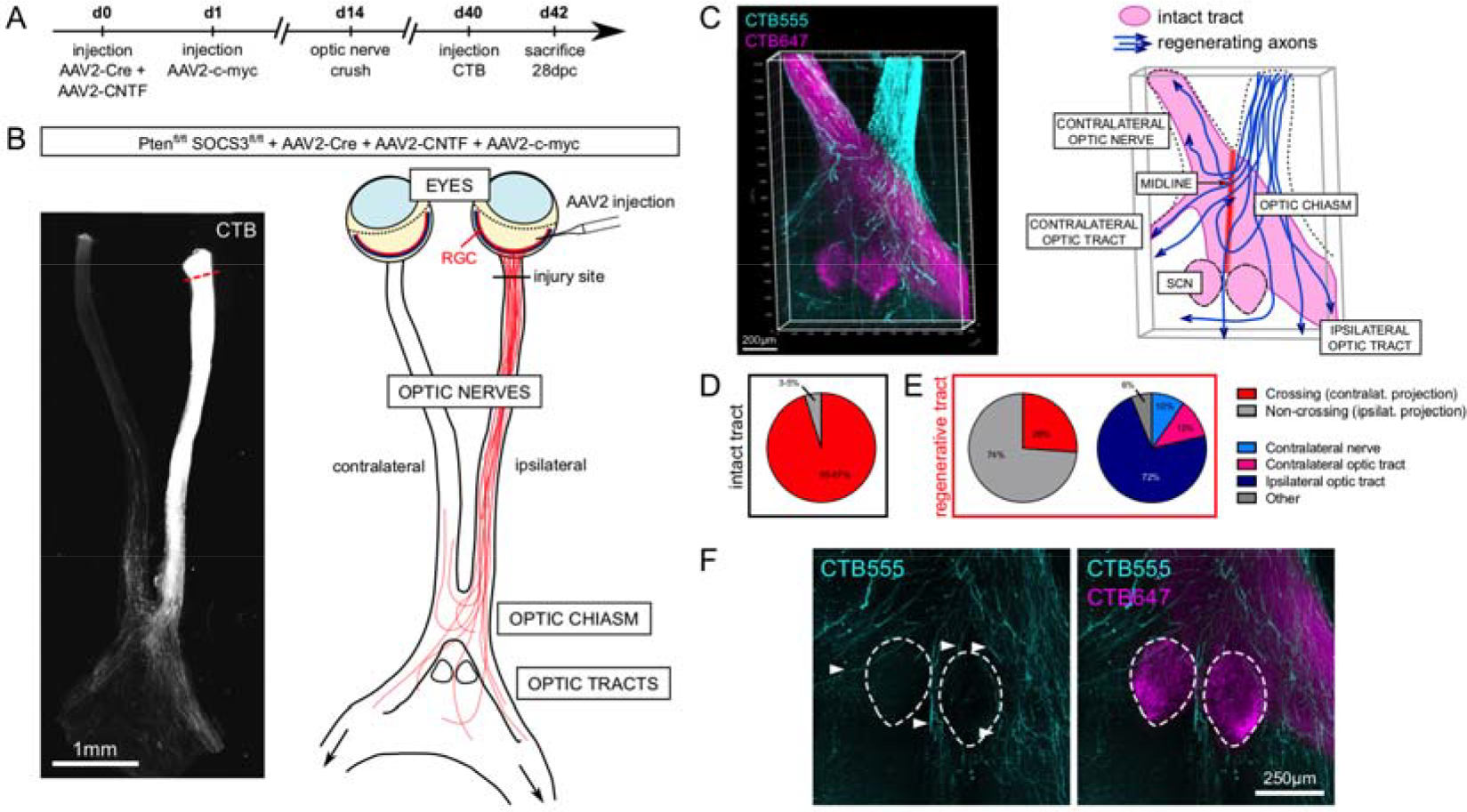
Regenerating axons fail to reform the visual circuit. (A) Timeline of experimental model of long-distance regeneration in the mouse visual system. (B) Confocal picture of whole optic nerves and optic chiasm 28 days post-unilateral optic nerve crush (28dpc) in one Pten^fl/fl^ SOCS3^fl/fl^ mouse eye injected with AAV2-Cre + AAV2-CNTF + AAV2-c-myc (AAV2-Cre/CNTF/c-myc). Regenerating axons are traced using CTB (white). The red dashed line indicates the lesion site. Scheme of guidance defects observed in unilateral optic nerve crush: some axons turn back to the ipsilateral eye, others project to the contralateral optic nerve or get lost in the optic chiasm. (C) 3D picture and scheme of whole optic nerves and optic chiasm 28dpc in unilateral optic nerve crush. Regenerating axons are traced with CTB555 (cyan). Intact (contralateral) axons are traced with CTB647 (magenta). (D) Proportion of midline crossing in intact circuit. (E) Quantification of guidance defects observed in long-distance regeneration model: regenerating axons fail to resume the ipsi-versus contralateral distribution of the intact circuit. Results are presented as the number of axons that cross the section of interest (e.g. midline, contralateral optic tract, ipsilateral optic tract) as a percentage of the total number of regenerating axons reaching the distal end of the ipsilateral optic nerve. (F) Confocal picture showing regenerating axons traced with CTB555 (cyan) and intact axons traced with CTB647 (magenta). Regenerating axons fail to enter the SCN.

For this purpose, we injured both optic nerves (bilateral ONC) and induced regeneration in only one eye (**Extended Data Figure 1A**). With this model, we observed similar axon pathfinding defects, but with a higher proportion of regenerating axons crossing the midline and invading the contralateral optic nerve and optic tract (**Extended Data Figure 1A-E**). However, regenerating axons still fail to enter the SCN (**Extended Data Figure 1F**). Thus, the presence of intact projections is not the cause of SCN reinnervation failure.

The SCN is normally innervated by the M1-intrinsically photosensitive RGC (ipRGC), which express high levels of the melanopsin photopigment ^18,19^. We analyzed ipRGC survival based on melanopsin expression. In the long-distance regeneration model, almost 95% of melanopsin^+^ RGC survive at 28dpc (**Extended Data Figure 2A**), consistent with previous studies ^20,21^. In addition, using a reporter mouse line expressing GFP under the melanopsin promoter (*Opn4*-GFP) ^22^, we found that ipRGC axons grow over long distances along the optic nerve (**Extended Data Figure 2B**). To confirm that melanopsin^+^ RGC axons reach the distal part of the optic nerve, we injected fluorescently-labeled cholera toxin B (CTB), a retrograde tracer, specifically in the optic chiasm (**Extended Data Figure 2C**). We could detect CTB^+^ melanopsin^+^ RGC in the retina (**Extended Data Figure 2D**). These results further confirm that SCN reinnervation failure is not due to an impaired survival and/or regeneration of ipRGC subpopulation, but rather that SCN itself has an intrinsic repulsive activity.

To characterize this repulsive activity, we conducted a candidate screen among canonical guidance cues ^23^ expressed in adult wild-type SCN (**Extended Data Figure 3A**). In addition to cues identified in our previous proteomics screen, such as NrCAM or PlexinA1 ^15^, we analyzed the expression of guidance molecules critical for visual system development, particularly in the developing optic chiasm ^24^. Interestingly, all members of the Slit family (*Slit1, Slit2* and *Slit3*) are expressed in the intact adult SCN. Their expression remains stable 28 days after bilateral ONC (**Extended Data Figure 3A-B**). We confirmed that *Slit* mRNAs are still expressed in the SCN in the long-distance regeneration model (**Figure 2A**). This suggests that Slit guidance cues are good candidates. First, Slits are known to repel RGC axons through Robo1 and Robo2 receptors ^25^. Second, Slit and Robo knockouts display similar guidance defects of visual axons as regenerating RGC axons in our regenerative model, with disorganized growth at the optic chiasm and projection into the contralateral optic nerve ^26^. Thus, we hypothesized that Slits expressed in the SCN could block the reinnervation of SCN neurons by Robo-expressing RGC axons. We first analyzed *Robo1* and *Robo2* expression in mature intact and injured RGC. We showed that *Robo1* and *Robo2* mRNAs are expressed in RGC and remain high at 28dpc (**Extended Data Figure 3C**). In particular, melanopsin^+^ RGC express *Robo1* and *Robo2* (**Extended Data Figure 3D**).

**Figure 2:**
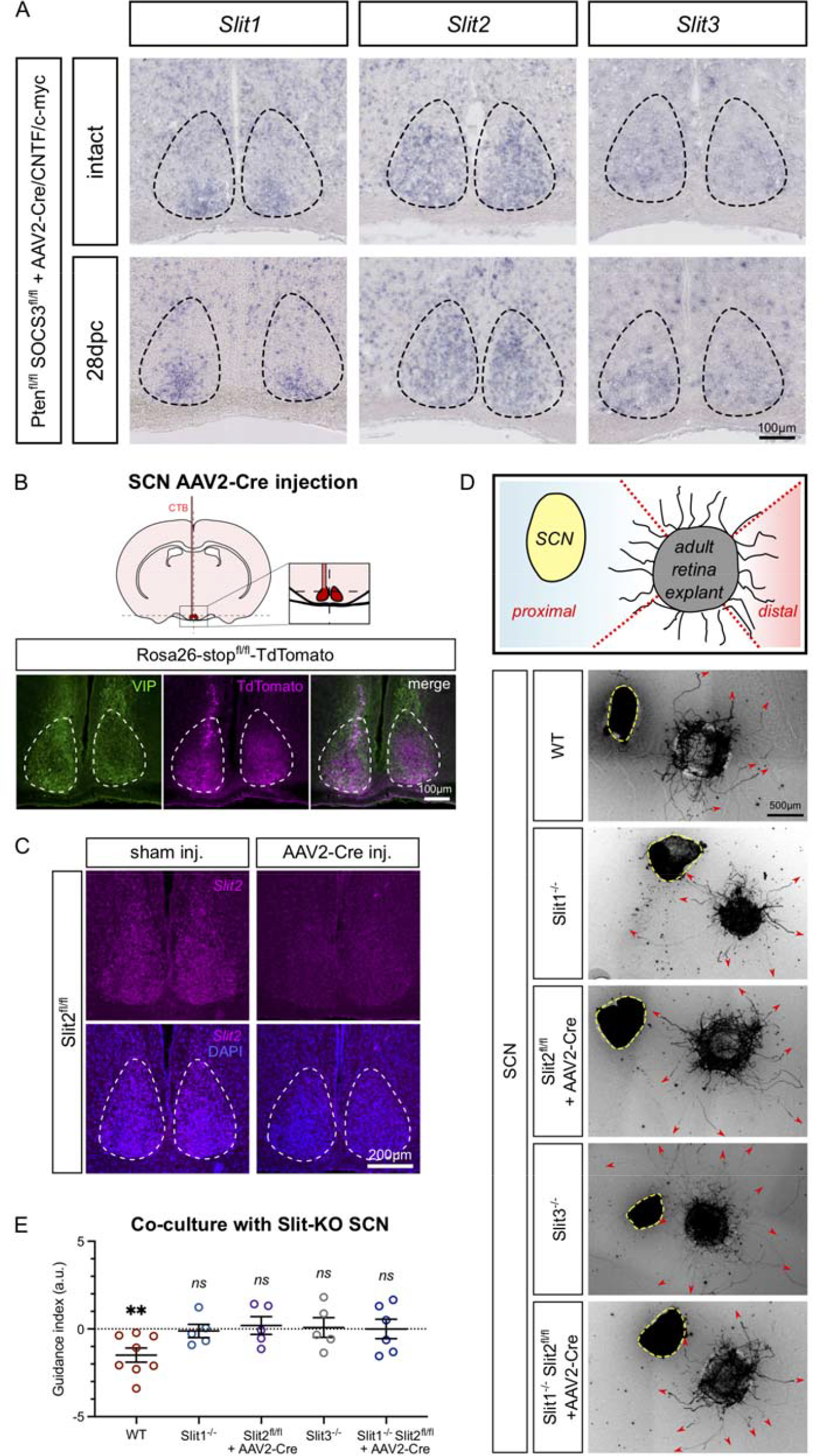
The SCN exerts a Slit-dependent repulsive activity. (A) In situ hybridization showing *Slit1, Slit2* and *Slit3* expression in the SCN of Pten^fl/fl^ SOCS3^fl/fl^ mice, where one eye was injected with AAV2-Cre/CNTF/c-myc. Expression is maintained at 28dpc. (B) Principle of stereotaxic injection of AAV2-Cre in the SCN. Control of injection in the SCN of Rosa26-stop^fl/fl^-TdTomato mice showing TdTomato expression (magenta) in the SCN labeled with anti-VIP antibody (green). (C) In situ hybridization showing *Slit2* deletion in Cre-injected SCN of a Slit2^fl/fl^ mouse. (D) The WT SCN exerts a repulsive activity on axons growing from a retina explant, whereas depletion of Slit1, Slit2 or both from the SCN abolishes its repulsive activity on growing axons ex vivo. Representative pictures of a WT, Slit1^-/-^, Slit2^fl/fl^ + AAV2-Cre, Slit3^-/-^ or Slit1^-/-^ Slit2^fl/fl^ + AAV2-Cre SCN, co-cultured with a retina explant from a Pten^fl/fl^ SOCS3^fl/fl^ eye injected with AAV2-Cre/CNTF/c-myc. (E) Corresponding quantification of guidance index showing the log-ratio of axon growth in the proximal versus the distal area. A negative guidance index indicates a significantly repulsive activity, while a null guidance index indicates a neutral activity. Data are expressed as mean +/- s.e.m. One-sample t-test compared to zero, ** p = 0.0079, ns: not significant.

Additionally, we used *ex vivo* cultures of adult retina explants from the long-distance regeneration model ^27^ to show that Robo1 and Robo2 proteins are expressed on growth cones of regenerating axons (**Extended Data Figure 3E-F**). This suggests that regenerating axons, including ipRGC, can respond to Slits repellents.

Next, we co-cultured retina explants from the long-distance regeneration model and adult SCN explants. As observed *in vivo* (**Figures 1F** and **Extended Data Figure 1G**), wild-type (Slit-expressing) SCN exerts a repulsive activity on regenerating axons ex vivo (**Figure 2D**). To investigate whether this repulsive activity is mediated by Slits, we manipulated Slit expression in the SCN and analyzed RGC axons behavior. For this purpose, we used Slit1 and Slit3 full knockout mice. As Slit2 knockout is embryonic lethal ^28^, we induced Slit2 specific knockdown via targeted injection of AAV2-Cre in the SCN of Slit2^fl/fl^ mice. Cre expression was validated with a TdTomato reporter mouse line (**Figure 2B-C**). A significant loss of SCN repulsive activity was observed with SCN explants from Slit1, Slit2 and Slit3 single knockouts or Slit1/Slit2 double knockouts. In all cases, RGC axons grow in any direction including towards the Slit-deprived SCN (**Figure 2D-E**). We concluded that Slit family members contribute to SCN repulsive activity and may account for its reinnervation failure *in vivo*.

To decrease Slit repulsion *in vivo*, we deleted Robo1 and Robo2 receptors in adult RGCs prior to ONC. In a wild-type background, Robo1/2 deletion did not induce any significant axon regeneration (**Extended Data Figure 4A**). Hence, Robo themselves do not have an intrinsic regenerative effect on RGC axons. Next, we crossed *Pten*^*fl/fl*^ *SOCS3*^*fl/fl*^ mice with *Robo1*^*-/-*^ and/or *Robo2*^*fl/fl*^ mice to modulate each or both receptor(s) in combination with long-distance regeneration of RGC. The inactivation of Robo1 had no significant effect on SCN innervation (**Extended Data Figure 4B-C**), while Robo2 deletion led to a modest increase in the number of regenerating axons entering the SCN (**Extended Data Figure 4D-E**). Strikingly, the simultaneous inactivation of Robo1 and Robo2 resulted in a robust and significant increase of the number of regenerating axons entering both SCN, and notably the ipsilateral one (**Figure 3A-C, Supplementary Movies S1-2**). In order to determine whether regenerating ipRGC could enter the SCN, we crossed the Pten^fl/fl^ SOCS3^fl/fl^ OPN4-GFP mouse line with Robo1^-/-^ Robo2^fl/fl^ mouse line. We detected CTB^+^ GFP^+^ fibers in the SCN (**Figure 3D**), showing that regenerating ipRGC are able to enter the SCN upon Robo1/2 modulation.

**Figure 3:**
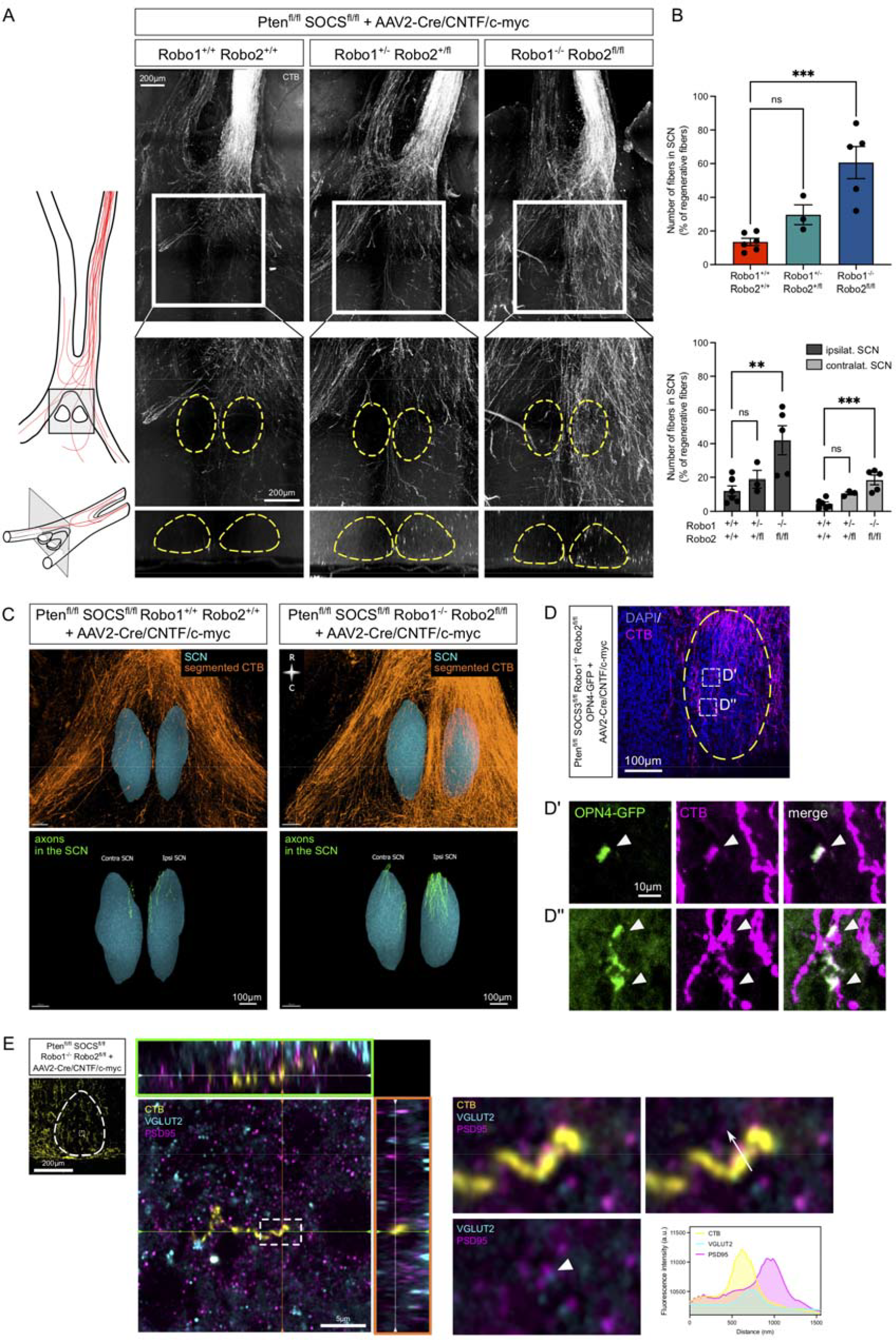
Modulation of Slit/Robo signaling in vivo promotes entry of regenerating axons into the SCN. (A) Confocal picture of whole optic nerves and optic chiasm at 28dpc in Pten^fl/fl^ SOCS3^fl/fl^ (left, Robo1^+/+^ Robo2^+/+^), Pten^fl/fl^ SOCS3^fl/fl^ Robo1^+/-^ Robo2^+/fl^ (centre) and Pten^fl/fl^ SOCS3^fl/fl^ Robo1^-/-^ Robo2^fl/fl^ (right) mice, with unilateral injection of AAV2-Cre/CNTF/c-myc and bilateral optic nerve crush. Regenerating axons are traced using CTB (white). Pictures are representive of N=3-6 animals in each condition. Zoom pictures: the SCN is indicated with the yellow dashed line on the maximum projection panel and on the XZ orthogonal section panel. (B) Top: quantification of the number of axons entering the SCN. Data are expressed as mean +/- s.e.m. One-way ANOVA with Dunnett’s correction, *** p-value = 0.0004, ns: not significant. Bottom: distribution of axons entering the ipsilateral and the contralateral SCN. Data are expressed as mean +/- s.e.m. One-way ANOVA with Dunnett’s correction, ** p-value = 0.0065, *** p-value = 0.0003, ns: not significant. (C) 3D imaging and reconstruction of regenerating axons in the SCN region, in Pten^fl/fl^ SOCS3^fl/fl^ (left) or Pten^fl/fl^ SOCS3^fl/fl^ Robo1^-/-^ Robo2^fl/fl^ (right) mice with unilateral injection of AAV2-Cre/CNTF/c-myc and bilateral optic nerve crush. (D) Confocal pictures of the ipsilateral SCN from a Pten^fl/fl^ SOCS3^fl/fl^ Robo1^-/-^ Robo2^fl/fl^ OPN4-GFP mouse injected with AAV2-Cre/CNTF/c-myc with bilateral optic nerve crush at 28dpc. Arrowheads point to GFP^+^ CTB^+^ fibers in the SCN, indicating that OPN4-GFP^+^ ipRGC enter the SCN in Robo-knockout condition at 28dpc. (E) Regenerating axons for synapses with target cells in the SCN. Left: confocal picture of the ipsilateral SCN from a Pten^fl/fl^ SOCS3^fl/fl^ Robo1^-/-^ Robo2^fl/fl^ mouse injected with AAV2-Cre/CNTF/c-myc with bilateral optic nerve crush at 28dpc. Centre: 3D confocal picture with Airyscan technology of an individual regenerative fiber forming a synapse in the SCN. The pre-synaptic compartment marker VGLUT2 (cyan) colocalizes with the CTB^+^ fiber (yellow) and is adjacent to the post-synaptic compartment marker PSD95 (magenta). Right: zoom showing the synapse formed by the CTB^+^ regenerative fiber. Profile of intensity in a single z plan along the white arrow.

To evaluate neuronal circuit reformation, we asked whether SCN-entering regenerating axons could form synapses in the SCN. For this purpose, we stained the SCN for pre-synaptic marker VGLUT2, expressed in ipRGC-SCN glutamatergic synapses ^29,30^, and post-synaptic marker PSD95. Using high-resolution imaging, we found VGLUT2^+^ synapses in CTB^+^ fibers that were directly adjacent to a PSD95^+^ post-synaptic compartment, indicative of synapses formed by regenerating fibers entering the SCN (**Figure 3E**). As synapses were found in the SCN, we then asked whether the newly formed neuronal circuit was functional in response to direct light activation. For this purpose, we set up a neuronal activity assay, based on expression of the immediate early gene c-fos in response to a short light exposure (**Figure 4A**). c-fos expression is triggered specifically in the SCN of wild-type intact mice exposed to light for 30 minutes in the night phase, both in terms of number of activated neurons and of c-fos intensity (**Extended Data Figure 5A**). In contrast, c-fos expression is not detected in non-photo-induced animals, nor upon bilateral ONC (**Extended Data Figure 5A**). In the long-distance regeneration model, we compared c-fos activation upon Robo1 and Robo2 modulation (*Robo1/2*^*-/-*^ and *Robo1/2*^*+/-*^) versus control (*Robo1/2*^*+/+*^). We found a significant increase in the number of c-fos^+^ cells in the SCN, as well as a significant increase of c-fos intensity (**Figure 4B**). Since *Robo2* knockout also induced a modest SCN innervation (**Extended Data Figure 4D-E**), we assessed c-fos expression in *Robo2*^*-/-*^ condition, and observed a significant increase in c-fos intensity compared to *Robo2*^*+/+*^ control condition (**Extended Data Figure 5B**). Altogether, these results strongly suggest that SCN reinnervation by regenerating axons leads to specific activation in target cells upon light exposure (**Extended Data Figure 5C**).

**Figure 4:**
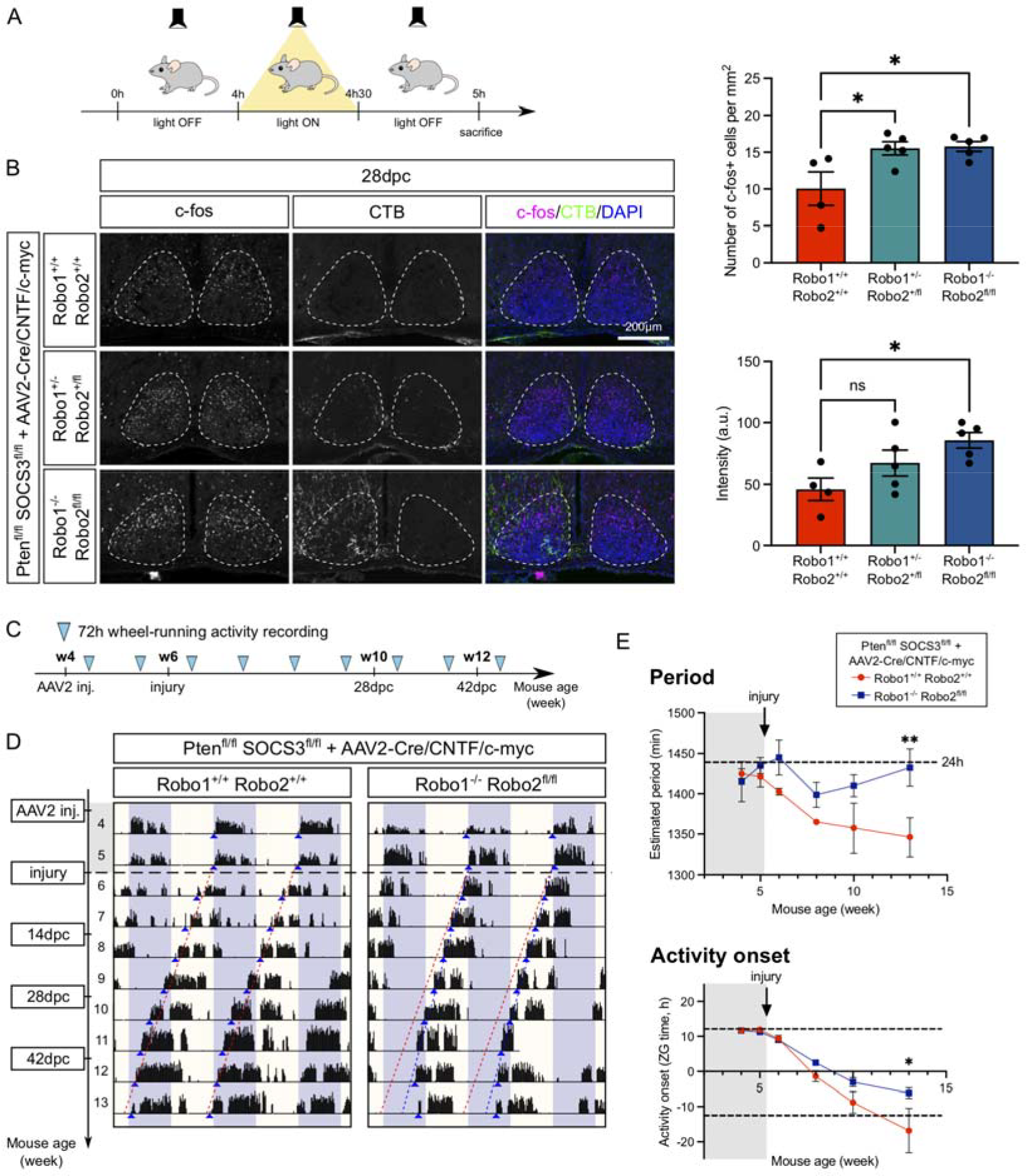
Guidance of regenerating axons into the SCN results in its reactivation. (A) Principle of c-fos activation experiment. (B) Confocal pictures showing SCN expression of c-fos (magenta) in the 28dpc regeneration model, in Robo1^+/+^ Robo2^+/+^, Robo1^+/-^ Robo2^+/fl^ and Robo1^-/-^ Robo2^fl/fl^ conditions. CTB (green) was used to trace regenerating axons. The SCN is indicated with a white dashed line. Each picture is representative of N=5 mice. Corresponding quantification of c-fos^+^ cell number per mm^2^ section in the SCN (top) and of c-fos intensity in c-fos^+^ cells in the SCN (bottom). Data are expressed as mean +/- s.e.m. One-way ANOVA with Dunnett’s correction, * p-value < 0.05, ns: not significant. (C) Timeline of wheel-running assay in the long-distance regeneration model. (D) Representative 72h actograms recorded every week in relation to environmental light/dark cycle, in two conditions: Pten^fl/fl^ SOCS3^fl/fl^ Robo1^+/+^ Robo2^+/+^ (left) and Pten^fl/fl^ SOCS3^fl/fl^ Robo1^-/-^ Robo2^fl/fl^ (right). Blue arrowheads point to activity onset. Activity onset slope is shown by the red dashed line in Robo1^+/+^ Robo2^+/+^ condition and the blue dashed line in Robo1^-/-^ Robo2^fl/fl^ condition. (E) Top: quantification of the estimated period in minutes calculated using the Chi-2 periodogram method. Data are expressed as mean +/- s.e.m. Two-way ANOVA with Sidak’s correction, ** p-value = 0.0079. Bottom: quantification of activity onset in Zeitgeber hours (hours to beginning of period, light on). Data are expressed as mean +/- s.e.m. Two-way ANOVA with Sidak’s correction, * p-value = 0.0201.

To further challenge this observation, we performed two additional experiments. First, we asked whether activated SCN neurons were functionally connected by regenerating axons. To this end, we stained the SCN innervated by Robo1^-/-^/Robo2^fl/fl^ regenerating axons for synaptic markers and c-fos. Using high-resolution microscopy (Airyscan confocal), we detected the presence of synapses (VGLUT2^+^/PSD95^+^ adjacent compartments) from CTB^+^ fibers and activated c-fos in SCN target cells (**Extended Data Figure 6A**). Second, we asked whether c-fos expression correlates with CTB^+^ regenerating axons distribution in the SCN. We performed an automatic detection of c-fos^+^ and CTB^+^ puncta combined with spatial point pattern analysis ^31^. We then plotted the density maps of c-fos and CTB across reinnervated SCN sections. The distribution of distances between CTB^+^ and c-fos^+^ puncta shows that c-fos activation clusters where CTB^+^ regenerating fibers enter in the SCN (**Extended Data Figure 6B-C**). This is the case in intact condition, with a detection of both types of events in the SCN core region (**Extended Data Figure 6B**), consistent with the facts that, during development, RGC axons enter the SCN ventrally ^32^ and that connections are more clustered in the SCN core. In the regenerative condition, c-fos is activated mostly in the SCN shell region, and this activation coincides with the dorsal entry point of CTB^+^ regenerating fibers in the SCN (**Extended Data Figure 6C**). Comparison of c-fos^+^ distributions shows that c-fos is more activated in the SCN core in intact condition while during regeneration c-fos is more activated in the shell (**Extended Data Figure 6D)**. Altogether, these results indicate that the newly formed retinorecipient circuit triggers functional neuronal activation in the SCN.

Finally, we asked whether this neuronal circuit could restore function. For this purpose, we took advantage of the SCN role as the master regulator of circadian rhythms to assess circuit recovery at the behavioral level. By monitoring running wheel activity, we compared two groups of animals: *Pten*^*fl/fl*^ *SOCS3*^*fl/fl*^ + AAV2-Cre/CNTF/c-myc (long-distance regeneration with no SCN reinnervation) and *Pten*^*fl/fl*^ *SOCS3*^*fl/fl*^ *Robo1*^*-/-*^ *Robo2*^*fl/fl*^ + AAV2-Cre/CNTF/c-myc (long-distance regeneration with SCN reinnervation and activation) (**Figure 4C**). At the time of AAV2 injection, both groups show similar circadian rhythms, with a clear activity at night (dark phase) and sleep at day (light phase), and a circadian period indistinguishable from 24h (1440min), imposed by the light/dark shift in the environment (**Figure 4D-E**). At the time of bilateral ONC, mice lose visual information relay and adopt a spontaneous circadian rhythm shorter than 24h (**Figure 4D**), as described for animals kept continuously in the dark (no light stimulation) ^33^. From this time on, both groups undergo a continuous earlier onset activity due to shorter intrinsic period, up to 14dpc. However, we observed that this forwarded onset activity is smaller from 28dpc in the reinnervated SCN group, and is significantly reduced from 42dpc (**Figure 4D-E**), with a period going back to 24h (**Figure 4E, Extended Data Figure 7**). Although the light/dark cycle is not fully reestablished, these results suggest that the reinnervated SCN group responds to an environmental 24h-light/dark period, unlike the control group that continuously adopts its intrinsic circadian period regardless of environmental visual input. Therefore, SCN reinnervation upon axon guidance restores a functional circuit.

Altogether, our study demonstrates that axon guidance is essential for visual function recovery during optic nerve regeneration. Indeed, we show that: (i) the mature SCN expresses repulsive guidance cues that counteract its reinnervation; (ii) modulation of the repulsive signaling makes regenerating axons enter the SCN; and (iii) this reinnervation is associated with synapse formation, target cell activation and functional recovery. So far, promoting axon regeneration/outgrowth failed to achieve full functional recovery. One reason is that across all regenerative models, severe axon pathfinding defects lead to little or no target reinnervation ^13^. These observations led us to reevaluate the developmental process of axon guidance in a mature circuit, which represents one of the main challenges in CNS repair. Previously, guidance cues have been studied in the context of regeneration, but only at the lesion site. Indeed, upon injury there is an upregulation of repulsive guidance factors associated with regeneration failure ^34–36^. Modulation of these cues was mainly used to assess their role in axonal growth, but not their guidance properties ^37,38^.

Axon guidance mechanisms are dynamically regulated during development, shaping precise neuronal connectivity and circuit formation. In the context of CNS repair, resuming correct axon trajectories relies on the fine spatial characterization of the mature CNS environment, which is very different in nature and distribution than the embryonic one. Recently, we adopted an unbiased, proteomics-based approach to establish a map of guidance cues and associated factors in RGC primary brain targets of the mature visual system ^15^. This mature guidance landscape is durably expressed in adulthood and remains stable even upon bilateral ONC. Therefore, triggering axon regrowth is not sufficient to form a neuronal circuit as regenerating axons face a refractory environment for reinnervation. Yet, manipulation of associated receptors in RGC leads to re-routing of axons, showing that regenerating axons have the potential to be guided ^15^.

In the present study, the inhibition of Slit/Robo signaling provides the first proof-of-concept that targeted functional reinnervation of the brain by regenerating axons is regulated by axon guidance cues. Furthermore, our results show that mature SCN innervation is different from the developmental stage. Indeed, most regenerating axons enter the SCN dorsally, consistent with neuronal activation in the SCN shell, whereas during development, ipRGC axons enter the SCN ventrally and connect mostly to the core region ^32^. These observations suggest that other mechanisms may influence SCN reinnervation, e.g. other guidance cues ^15^ or differential mechanical stiffness in the environment ^39^.

As during development, modalities of circuit repair may depend on the regeneration and guidance properties of each neuronal subpopulation. In this regards, several high-throughput data analyses have characterized the injury response and differential resilience of the multiple RGC subpopulations ^21,40–42^. In particular, in our case, ipRGC robust survival and regeneration, even late after injury, is consistent with other models of regeneration ^4,20,43^. Whether response to guidance cues is uniform across these different subpopulations remains unknown. Our experiments show a repulsive activity of Slit family members on regenerating axons, similar to embryonic neurons ^25,44^. However, depending on guidance cues, mature neuron subpopulations may actually differ both in expression of guidance receptors and in the guidance response itself. Therefore, our study opens a new field of research to decipher guidance mechanisms for specific neuronal circuits.

We show that circuit reorganization is associated with functional recovery at multiple levels: synapse formation, post-synaptic neuronal activation and behavioral recovery. We also uncovered unexpected phenotypes, notably an asymmetrical SCN reinnervation with the ipsilateral nucleus preferentially targeted. Interestingly, during development, each SCN is symmetrically connected to both eyes, unlike other retinorecipient brain nuclei such as the lateral geniculate nucleus and the superior colliculus. Indeed, almost 50% ipRGC project bilaterally to each SCN, via branching axons at the optic chiasm ^30^. In our regenerative model, it remains unclear whether these mechanisms are still at play. Nonetheless, we observed that both SCN respond symmetrically to photoactivation analyzed by c-fos expression. This may be due to the internal coupling that ensures correct synchronization of both SCN for circadian clock control ^45–47^.

Finally, our behavioral data show a significant recovery of the visual response to environmental light/dark cycles. This functional recovery does not fully resume the intact circuit, which could be due to mature SCN reinnervation properties. Indeed, the regenerating circuit preferentially activates the SCN shell, in contrast to the intact circuit that has preferential ipRGC projections to the core. Neurons from the shell and the core play different roles in the control of the circadian rhythms and other non-image forming behaviors ^16,48^, which may explain why functional recovery is incomplete in our model. In addition, ipRGC responses are mostly mediated by excitatory glutamatergic signaling ^29,49^, but ipRGC were recently shown to also release GABA in the SCN ^50^. This non-canonical inhibitory circuit mediates circadian photoentrainment in a light intensity-dependent manner ^50^. In the case of mature SCN reinnervation, how the newly formed circuit mediates visual information remains to be fully characterized.

To conclude, our results provide evidence that mature regenerating axons respond to external guidance cues, and that modulation of the relevant guidance signals can lead to functional neuronal circuit reconstruction in the adult brain. This study will set the basis of future therapeutic strategies to achieve circuit repair after CNS injury.

## Methods

### Animals

All animal care and procedures have been approved by the Ethics Committee of the Grenoble Institute of Neuroscience (project number 201612161701775) and by the French Ministry of Research (project number APAFIS#9145-201612161701775v3) in accordance with French and European guidelines. We used Pten^fl/fl^ SOCS3^fl/fl^, OPN4-GFP Pten^fl/fl^ SOCS3^fl/fl^, Pten^fl/fl^ SOCS3^fl/fl^ Robo1^-/-^ Robo2^fl/fl^, Pten^fl/fl^ SOCS3^fl/fl^ Robo1^-/-^, Pten^fl/fl^ SOCS3^fl/fl^ Robo2^fl/fl^, Slit1^-/-^, Slit2^fl/fl^, Slit1^-/-^ Slit2^fl/fl^ and Slit3^-/-^ mice in this study, regardless of their sex (except for behavior assay where only females were used) and aged at least 3 weeks. To obtain OPN4-GFP Pten^fl/fl^ SOCS3^fl/fl^ mice, we crossed OPN4-GFP mice ^22^ with Pten^fl/fl^ SOCS3^fl/fl 6^. To obtain Pten^fl/fl^ SOCS3^fl/fl^ Robo1^-/-^ Robo2^fl/fl^ mice, we crossed Pten^fl/fl^ SOCS3^fl/fl^ mice with Pten^fl/fl^ Robo1^-/-^ Robo2^fl/fl^ mice ^51,52^. To obtain OPN4-GFP Pten^fl/fl^ SOCS3^fl/fl^ Robo1^-/-^ Robo2^fl/fl^ mice, we crossed OPN4-GFP mice with Pten^fl/fl^ SOCS3^fl/fl^ Robo1^-/-^ Robo2^fl/fl^ mice. Mice were housed in standard housing conditions with a 12 h light/dark cycle. When possible, animals were housed in groups of 2–5 per cage. Animals were fed and watered ad libitum. Before each surgery, all animals were anesthetized with an intraperitoneal injection of ketamine (75-100 mg/kg, Clorkétam 1000, Vetoquinol) and xylasine (10 mg/kg, Rompun 2%, BAYER).

### Intravitreal virus injection

Intravitreal injections were performed as described ^27^. The eyelid of the eye was pinched with a mini bulldog serrefines clamp (Fine Science Tools) to bring out the eyeball and expose the posterior part of the eye. Using a Hamilton syringe (Hamilton) connected to a glass micropipette (Sutter Instruments), the posterior part of the eye located just behind the ora serrata was punctured with an angle of 45° to avoid damaging the lens. About 2μL of vitreous humor was taken out before injecting 1μL of adeno-associated virus type 2 (AAV2), concentrated with at least 10^11^ viral particles per mL. Viral vectors used in this study are AAV2-Cre, AAV2-CNTF and AAV2-c-myc. 2 days before sacrifice, animals received an intravitreal injection of 1μL of CTB at 1μg/μL (ThermoFisher Scientific). Mice with eye inflammation, damage or atrophies were excluded from further experiments.

### Optic nerve crush

2 weeks after the AAV2 intravitreal injections, optic nerve crush was performed. The connective tissue above the eye was cut using scissors (Fine Science Tools). Forceps (Fine Science Tools, 11251-20) were then placed between the 2 arteries located behind the eyeball in order to expose the optic nerve. Using a second pair of forceps, the optic nerve was pinched for 5 seconds at 1-2 mm behind the eyeball. Animals with excessive bleeding were excluded from further experiments.

### Intracranial injections

10-week-old Pten^fl/fl^ SOCS3^fl/fl^, Slit2^fl/fl^ or Slit1^-/-^ Slit2^fl/fl^ mice were anesthesized and installed on a stereotaxic frame. The skin was cut letting the skull visible. Using the canula, the bregma was defined as the origin of axes. After verifying flat skull positioning, one hole was performed using a dental burr at A/P: +1mm, M/L: +0.1mm, D/V: -6mm (for SCN injection) and -6.5mm (for chiasm injection). 1µL of virus (AAV2-Cre for co-culture experiment, Slit2^fl/fl^ or Slit1^-/-^ Slit2^fl/fl^ mice) or CTB (for retrograde labeling, Pten^fl/fl^ SOCS3^fl/fl^) was injected using a syringe pump (0.5µL/min). After injection, the skin was sutured and animals were sacrificed 2 weeks later.

### Neuronal activation assay

To study SCN neuron activation upon light exposure, 4h after the dark cycle start, we exposed mice for 30min to light before going back 30min to dark (4h dark - 30min light - 30min dark; this scheme is called 4-30-30). We designed 5 mice groups. Group 1 (positive control): WT intact mice with 4-30-30 condition. Group 2 (negative control): WT intact mice kept in the dark. Group 3: WT mice with bilateral optic nerve lesion in 4-30-30 condition. Group 4: Pten^fl/fl^ SOCS3^fl/fl^ Robo1^+/+^ Robo2^+/+^ mice in 4-30-30 condition. Group 5: Pten^fl/fl^ SOCS3^fl/fl^ Robo1^+/-^ Robo2^+/fl^ mice in 4-30-30 condition. Group 6: Pten^fl/fl^ SOCS3^fl/fl^ Robo1^-/-^ Robo2^fl/fl^ in 4-30-30 condition. Groups 4 to 6 received a unilateral injection of AAV2-Cre/CNTF/c-myc as described above, then a bilateral ONC two weeks later. Two days before sacrifice, groups 4 to 6 received an intravitreal injection of CTB to control for complete injuries. For groups 4-6, neuronal activation assay was performed at 28dpc. After 4-30-30 conditioning, mice were sacrificed and perfused in the dark. Brains were dissected out and sectioned into 40µm thick-section. Immunofluorescence with anti-c-fos antibody was performed.

### Adult retina explant cultures

Adult retina explant cultures were performed according to ^27^. Both eyes of 4-week-old Pten^fl/fl^ SOCS3^fl/fl^ mice were injected with AAV2-Cre + AAV2-CNTF on day 0, then AAV2-c-myc on day 1. Two weeks later, mice were sacrificed by cervical dislocation and retinas were dissected out in Hibernate-A and cut into small pieces (about 500μm in diameter, as described ^27^). Explants were laid on glass coverslips previously coated with 0.5mg/mL poly-L-lysine overnight at room temperature, then with 20µg/mL laminin for 2h at room temperature, then with a thin layer of coating medium (4mg/mL methylcellulose, 2% B27 (ThermoFisher Scientific), 1% L-Glutamine (Corning) in Hibernate-A). Explants were cultured for 7 to 10 days before fixation and immunofluorescence.

### Co-culture experiments

Two weeks after intracranial injection of Slit2^fl/fl^ or Slit1^-/-^ Slit2^fl/fl^ mice, animals were sacrificed by cervical dislocation. Fresh brains were sectioned using a vibratome (300µm thick sections). The two SCN were microdissected out, each constituting one SCN explant. Glass coverslips used for the co-cultures were previously coated with 0.5mg/mL poly-L-lysine for 1h at 37°C. After washes and air-dry, a thin layer of collagen matrix was applied on each coverslip (500µL of 3mg/mL collagen I (from rat tail, Thermo Scientific), 20µL of 1mg/mL laminin (Sigma-Aldrich), 12.5µL of 1N NaOH, 100µL of PBS 10X (Euromedex) and 367.5µL of Neurobasal-A) and left to polymerise at 37°C for 20 min. In parallel, retina from Pten^fl/fl^ SOCS3^fl/fl^ previously injected with AAV2-Cre/CNTF/c-myc were dissected out and prepared for explants. A retina explant and a SCN explant were laid onto the coated coverslip and a 60µL drop of the collagen mix was applied. Co-cultures were placed at 37°C for few minutes before adding the culture medium: 2% B27, 1% L-Glutamine, 1% Penicillin/Streptomycin (ThermoFisher Scientific) in Neurobasal-A (ThermoFisher Scientific). Co-cultures were incubated at 37°C with 5% CO_2_ for 10 days before fixation and immunofluorescence.

### Tissue preparation

At 28dpc animals were anesthetized with ketamine (75-100 mg/kg) and xylasine (10 mg/kg) and then intracardially perfused with ice-cold PBS and ice-cold 4% paraformaldehyde dissolved in PBS (PFA) (Electron Microscopy Sciences). Brains, optic nerves and eyes were dissected and post-fixed in 4% PFA overnight at 4°C. Brains and eyes were dehydrated in 30% sucrose (Sigma-Aldrich) for 2 days at 4°C before embedding in OCT tissue freezing medium (MM-France) and stored at -80°C. Using cryostat sectioning, 20µm coronal brain, 14µm eyes and 14µm optic nerve sections were performed. Sections were collected on SuperFrost slides (ThermoFisher Scientific) and stored at -20°C until further use.

### Optic nerve and chiasm clearing

Optic nerves, chiasm and/or SCN were clarified as described ^27^. First, samples were post-fixed in 4% PFA overnight at 4°C. If necessary, samples were first permeabilized in PBS 0.5% Triton X-100 for 2h at room temperature and incubated in 10µg/mL DAPI (Sigma-Aldrich-Merck, D9542) for 30min, then washed several times in PBS 0.5% Triton X-100. Alternatively, samples were permeabilized, blocked in PBS 0.5% Triton X-100 5% donkey serum and stained for anti-VIP (1:400, Immunostar) overnight at 4°C. Secondary antibody incubation was performed for 2h at room temperature, then samples were washed several times in PBS 0.5% Triton X-100. Next, samples were gradually dehydrated in ethanol baths (50%, 80%, 95%, 100%) for at least 20min each at room temperature. Samples were incubated in 100% ethanol overnight at 4°C. Samples were further incubated in hexane (Sigma-Aldrich-Merck) for 2h before clearing in benzyl alcohol:benzyl benzoate (1:2) (Sigma-Aldrich-Merck). Samples were stored in this solution at room temperature and protected from light.

### Immunostaining

For retina and brain sections, samples were first defrosted and washed three times 10min with PBS. Samples were saturated in a blocking solution (3% bovine serum albumin (BSA) (Sigma-Aldrich), 5% donkey serum (Merck Millipore), 0.5% Triton X-100 (Sigma-Aldrich) in PBS) for 1h at room temperature and then incubated overnight at 4°C with primary antibodies: anti-melanopsin (Abcam, 1:200), anti-GFP (Abcam, 1:500), anti-RBPMS (Merck Millipore, 1:400), anti-VIP (Immunostar, 1:400), anti-VGLUT2 (Abcam, 1:500), anti-PSD95 (Abcam, 1:500), anti-c-fos (Cell Signaling Technology, 1:200). The following day, three washes of 10 min with PBS 0.1% Triton X-100 were carried out before incubation for 2h with secondary antibodies diluted 1:500 in blocking solution: anti-chicken Alexa 488 (Jackson), anti-goat Alexa 488 (ThermoFisher Scientific), anti-rabbit Alexa 488 (ThermoFisher Scientific), anti-rabbit Alexa 568 (ThermoFisher Scientific), anti-rabbit Alexa 647 (ThermoFisher Scientific), anti-guinea pig Alexa 647 (Jackson), anti-mouse Alexa 647 (ThermoFisher Scientific), anti-mouse Alexa 568 (ThermoFisher Scientific). Sections were washed three times for 10min and then mounted with Fluoromount-G with DAPI mounting medium (ThermoFisher Scientific).

Whole-mount retinas were first washes three times 10min with PBS, then blocked in OBS 0.5% Triton X-100 5% donkey serum for 1h at room temperature. Primary antibody anti-melanopsin (Abcam, 1:300) and anti-Tuj1 (Biolegend, 1:500) and secondary antibodies were incubated respectively overnight at 4°C and for 2h at room temperature. After three 10min-washes in PBS 0.1% Triton X-100, retinas were mounted with Fluoromount-G with DAPI (ThermoFisher Scientific).

For growth cone immunostaining from retina explant cultures, samples were fixed by adding an equal volume of 3% sucrose 8% PFA in culture medium, with 15min incubation. After three washes with PBS, explants were incubated overnight in PBS 3% BSA with primary antibody: anti-Robo1 (R&D Systems, 1:400) or anti-Robo2 (gift from Dr Alain Chédotal ^53^, 1:500), and anti-Tuj1 (Biolegend, 1:800). After three washes with PBS, secondary antibodies with Alexa 647-conjugated phalloidin (ThermoFisher Scientific, 1:500) were incubated for 2h. After PBS washes, explants were mounted with Fluoromount G with DAPI (ThermoFisher Scientific).

For co-cultures, samples were fixed by adding an equal volume of 3% sucrose 8% PFA in culture medium, with 15min incubation. Explants were washed three times in PBS and then permeabilized 10min in PBS 0.1% Triton X-100. Primary antibody (anti-Tuj1, Biolegend, 1:400) and secondary antibody were diluted in blocking solution (0.1% Triton X-100, 3% BSA, 5% donkey serum in PBS) and samples were incubated respectively for 2h and for 1h at room temperature. After PBS washes, samples were mounted with Fluoromount-G with DAPI (ThermoFisher Scientific).

### In Situ Hybridization

In situ hybridization on sections was performed as described ^54^. For chromogenic in situ hybridization (ISH), slices were incubated with digoxigenin (DIG)-labeled probes for *Ephb1, Netrin1, Nrcam, Plexina1, Robo1, Robo2, Sema6d, Sfrp1, Sfpr2, Shh, Slit1, Slit2, Slit3* or *Vegf* overnight at 65°C. For fluorescent ISH, samples were incubated with digoxigenin-labeling probes for *Robo1* or *Robo2*. The DIG-labeled probes were amplified with horseradish peroxidase using Cy3.5 tyramide signal amplification kit (PerkinElmer) for 5 to 10 min.

### Quantitative PCR

For sample collection, 6-week-old mice wild-type mice received a bilateral ONC. After 28 days, both SCN were dissected from injured (28dpc) and uninjured (intact) mice. N=4-5 mice were used in each group. SCN were dissociated in Trizol (Ambion Life technologies) for total RNA extraction. 100ng of total RNA were used for reverse transcription using SuperScript II (Invitrogen). mRNA levels were assessed by qPCR (Biorad) for *Slit1, Slit2*, and *Slit3*, and normalized to *Gapdh* levels.

### Imaging

Immunofluorescence and fluorescence in situ hybridization (ISH) on sections were imaged with DragonFly spinning disk confocal microscope from Andor, with a 25x objective. Chromogenic ISH on sections were imaged using Zeiss Slide Scanner Axio Scan.Z1. Co-cultures were imaged with epifluorescence microscope Nikon Ti Eclipse with a 4x objective. Robo immunofluorescence in growth cones and VGLUT2/PSD95 immunofluorescence in SCN were imaged using a confocal microscope (LSM710, Zeiss), with a 63x objective and AiryScan mode. Whole-cleared optic nerves, chiasm and SCN were imaged with DragonFly spinning disk confocal microscope from Andor, with a 20x objective. Images were acquired with 2µm-thick z-stack and were automatically stitched using Fusion software or with Imaris stitcher, then visualized and analyzed with Imaris software. CTB^+^ signal was segmented manually using Syglass software in virtual reality. SCN were segmented using DAPI^+^ signal on Imaris and fibers inside the SCN were segmented on Imaris. Movies were produced with Imaris and annotated with Adobe Premier Pro.

### Behavioral assay

For assessment of circadian rhythm, we monitored wheel-running activity of mice housed in a 12h light/dark cycle. Only females were considered in this study. Groups consisted in 4 Pten^fl/fl^ SOCS^fl/fl^ mice and 4 Pten^fl/fl^ SOCS^fl/fl^ Robo1^-/-^ Robo2^fl/fl^ mice. Each mouse was housed individually in a cage equipped with a connected wheel allowing to record spontaneous activity (Intellibio Innovation). Recordings were performed every week from the time of injection, and restricted to 72h to exclude surgeries, recovery time and litter changing. When not recorded, mice were still in presence of a running wheel. Data were acquired every 15min with ActiWheel software (Intellibio Innovation) and analysed with ActogramJ ^55^. For analysis, distance (in cm) was plotted versus time. The Chi-2 periodograms allowed to estimate the period for each recording session. Activity onset was assessed as the average time of starting activity across two or three 24h-recording periods. Recordings showing zero activity across a full period were excluded from analysis.

### Guidance defects quantification

Whole-cleared optic nerves, chiasm and SCN were imaged using DragonFly spinning disk confocal microscope (Andor) with a 20x objective. Images were acquired with 2µm-thick z-stack and were automatically stitched using Fusion software (20% tile overlap). All quantifications were assessed using Imaris software (version 9.6, Bitplane). For the percentage of regenerating axons into the contralateral optic nerve, we performed an XZ orthogonal section at the distal end of optic nerves using the ortho-slicer tool and we counted each CTB^+^ dot as one axon. We divided the axon number at the distal end of the contralateral optic nerve by the axon number counted at the distal end of the ipsilateral optic nerve (ION). For the proportion of regenerating axons in others regions, we proceeded in the same way, using the oblique-(YZ oblique section at the midline for the chiasm) or ortho-(XZ orthogonal section at the proximal part of the optic tracts) slicers tools and we divided the axon number by the axon number found at the distal end of the ION. The proportion of regenerating axons into the SCN was calculated using XZ orthogonal slicer tool. We counted the number of CTB^+^ fibers into each SCN and divided this value by the axon number present at the distal end of the ION.

### Quantification of melanopsin-positive neuron survival

Eyes from intact, 3 days post-crush (dpc), 14dpc and 28dpc Pten^fl/fl^ SOCS3^fl/fl^ mice (previously injected with AAV2-Cre/CNTF/c-myc) were collected and post-fixed in 4% PFA solution overnight at 4°C. Tissue were prepared and stained with anti-melanopsin and anti-RBPMS antibodies as described above. Using Zeiss Slide Scanner Axio Scan.Z1, retina slices were imaged with 20x objective. Melanopsin^+^ cells were counted on 5 retina sections for each animal and sum up. Using anti-RBPMS staining, we assessed the retina length of all retina sections. The sum of melanopsin^+^ cells number for each animal was normalized to the length of retina section (in cm). For each condition, the melanopsin^+^ cell survival was calculated by dividing the average melanopsin^+^ cell number of each animal by the average of intact condition.

### Co-culture data analysis

For each co-culture, the retina and the SCN explants were manually annotated using ImageJ and the proximal and distal regions were defined as the 90°-angle portion of retina explant towards and away from the SCN explant, respectively. Axon outgrowth was quantified in each region with a Sholl analysis using the ImageJ plug-in Neurite-J ^56^. Background noise filtering was performed automatically and manually corrected. The number of neurite intersects was determined by the Sholl analysis with a step of 25μm. The guidance index is calculated as the log2-transform of the total number of intersects in the proximal region versus the distal region. With this index, a positive value represents attraction towards the source (SCN explant), while a negative value represents repulsion away from the source, and a zero-value represents no preference. For statistical analysis, data were subjected to a one-sample t-test with a theoretical value of 0, using GraphPad Prism version 9.1.2.

### Spatial point pattern analysis

To correlate CTB and c-fos distributions in SCN sections, a spatial point pattern analysis was performed. The SCN was manually defined from the DAPI channel. Individual events were automatically detected from the immunofluorescence confocal images using Qupath ^57^. Multitype (CTB^+^ and c-fos^+^ events) point patterns were analysed using spatstat R package ^31^. Density maps with standard error intensity estimate were plotted. The G-cross function, i.e. the cumulative distribution of nearest neighbour distance from CTB^+^ events to c-fos^+^ events was plotted, taking into account the inhomogenous distribution of each marker with the function Gcross.inhom.

### Statistical analysis

All quantitative data are represented as mean +/- standard error of the mean (s.e.m.). All statistical analysis were performed using GraphPrism software version 8. Student’s t-test for two conditions or one-way ANOVA with Dunnett’s correction for at least three conditions were performed. For behavioral assay, Chi-2 periods and activity onset shift were compared with two-way ANOVA with Sidak’s correction. The difference between two averages were considered significant when the p-value was strictly below 0.05, with the following usage: * p-value < 0.05; ** p-value < 0.01; *** p-value < 0.001; and ns: p-value ≥ 0.05, not significant.

## Acknowledgements

We warmly thank Dr Alexandra Rebsam (Institut de la Vision, Paris, France) and Dr Julien Falk (INMG, Lyon, France) for their comments and feedbacks during Céline Delpech thesis committee meetings. We warmly thank Dr Xavier Nicol (Institut de la Vision, Paris, France), Dr Julien Courchet (INMG, Lyon, France) and Dr Mireille Albrieux (Grenoble Institute Neuroscience, France) for their comments and feedbacks during Noemie Vilallongue thesis committee meetings. We would like also thank Charlotte Corrao for *Slit1* ISH plasmid cloning. This work was supported by the Photonic Imaging Center of Grenoble Institute Neuroscience (Univ Grenoble Alpes – Inserm U1216) which is part of the IsdV core facility and certified by the IbiSA label. This work was supported by a grant from ANR (C7H-ANR16C49), from European Research Council (ERC-St17-759089) and NRJ foundation to HN. This work was supported by grants from the French National Research Agency in the framework of the “investissements d’Avenir” program (ANR-15-IDEX-02 NeuroCoG (HN, SB and CD)). JS is supported by Fondation pour la Recherche Médicale (FRM) postdoctoral fellowship (SPF201909009106). CD is supported by Fondation de France Bourse Berthe Fouassier. NV is supported by Fondation pour la Recherche Médicale (FRM) thesis fellowship (FDT202204014716).

## Author contributions

Conceptualization: HN. Methodology: HN. Investigation, formal analysis and validation: CD, JS and NV. Investigation: AB. Supervision: SB and HN. Writing – original draft: CD, JS, NV and HN. Investigation and resources: EP, FA, AP, BE. Funding acquisition: HN. Resources: AC, VC and YZ. Writing – review and editing: JS, NV, HN, AC and SB.

## Competing interests

The authors declare no competing interest.

## Material and correspondence

Data that support the findings of this study are either provided as supplementary material or are available from the lead contact upon request.

## Notes

### Competing Interest Statement

The authors have declared no competing interest.

