## Supplemental material for "Axon guidance during CNS regeneration is required for specific brain innervation"

**Extended data**

- Extended Data Figures 1 to 7
- Supplementary Movie captions
- Resource table for Materials and Methods

### Extended Data Figure 1

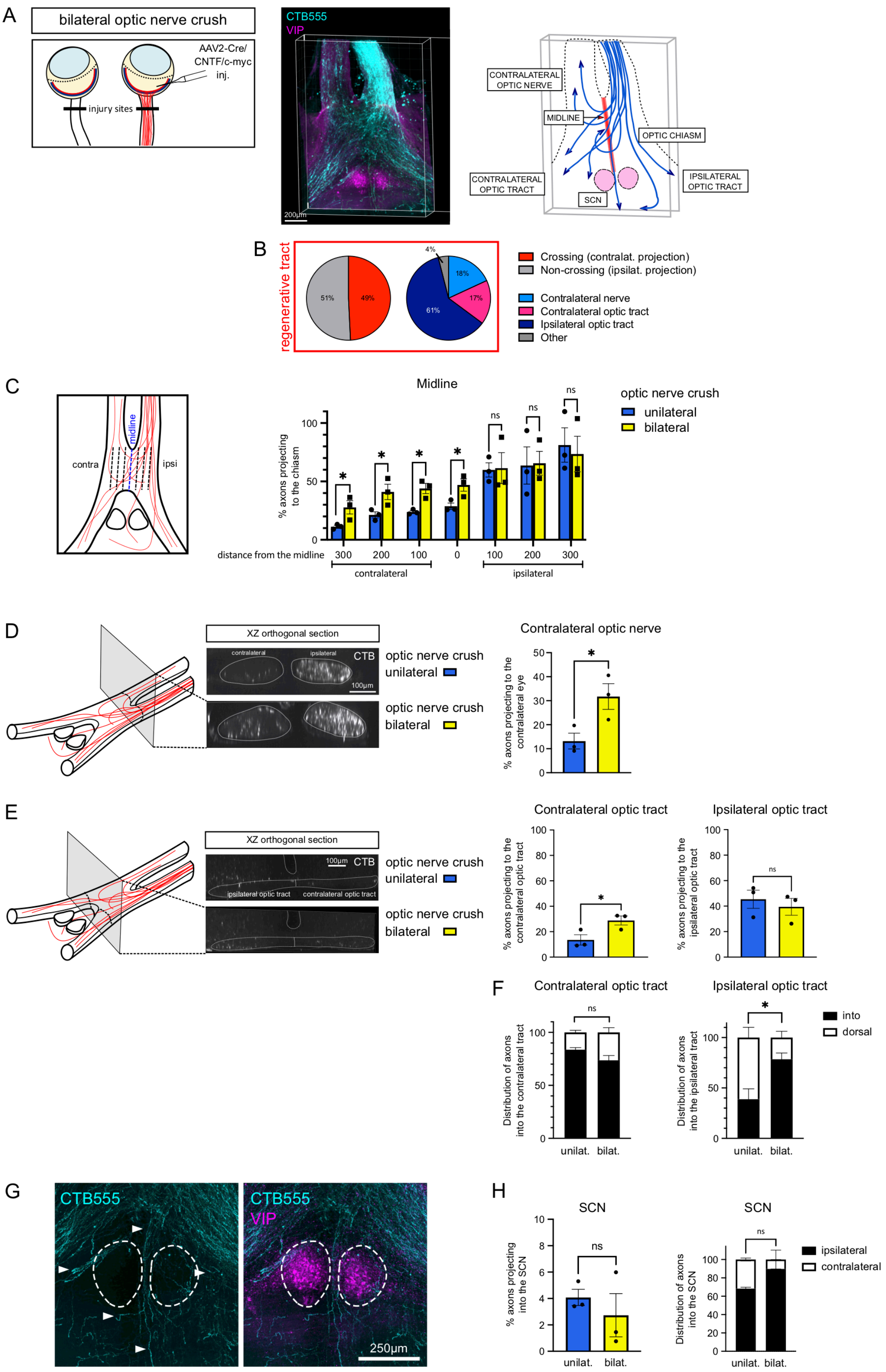


**Extended Data Figure 1: Regenerating axons display guidance defects in the adult visual system**. (A) Scheme of bilateral optic nerve crush. 3D picture and scheme of whole optic nerves and optic chiasm 28dpc in bilateral optic nerve crush. Regenerating axons are traced with CTB (cyan). The SCN is labeled with anti-VIP (magenta). (B) Quantification of guidance defects observed in bilateral optic nerve crush: regenerating axons fail to resume the ipsi- versus contralateral distribution of the intact circuit. Results are presented as the number of axons that cross the section of interest (e.g. midline, contralateral optic tract, ipsilateral optic tract) as a percentage of the total number of regenerating axons reaching the distal end of the ipsilateral optic nerve. (C) Quantification of the percentage of regenerative fibers intersecting the midline (distance 0) and projecting ipsilaterally or contralaterally in the optic chiasm. Comparison of unilateral and bilateral optic nerve crush. Unpaired t-tests, * p-value < 0.05, ns: not significant. (D) Representative confocal pictures of XZ orthogonal sections in the ipsi- and contralateral optic nerves. Quantification of the percentage of regenerative fibers projecting to the contralateral optic nerve. Comparison of unilateral and bilateral optic nerve crush. Data are expressed as mean +/- s.e.m. Unpaired t-test, * p-value < 0.05. (E) Representative confocal pictures of XZ orthogonal sections in the ipsi- and contralateral optic tracts. Quantification of the percentage of regenerative fibers projecting to the contralateral and ipsilateral optic tracts. Comparison of unilateral and bilateral optic nerve crush. Data are expressed as mean +/- s.e.m. Unpaired t-test, * p-value < 0.05, ns: not significant. (F) Distribution of axons projecting ipsi- or contralaterally, entering into the tract and dorsal to it. Comparison of unilateral and bilateral optic nerve crush. Data are expressed as mean +/- s.e.m. Unpaired t-test on axons projecting into the tract, * p-value < 0.05, ns: not significant. (G) Confocal picture showing regenerating axons traced with CTB (cyan) and the SCN labeled with anti-VIP (magenta). Regenerating axons fail to enter the SCN. (H) Left: quantification of the percentage of regenerative fibers entering the SCN. Comparison of unilateral and bilateral optic nerve crush. Data are expressed as mean +/- s.e.m. Unpaired t-test, ns: not significant. Right: distribution of axons entering the ipsi- versus contralateral SCN. Comparison of unilateral and bilateral optic nerve crush. Data are expressed as mean +/- s.e.m. Unpaired t-test on axons projecting in the ipsilateral SCN, ns: not significant.

### Extended Data Figure 2

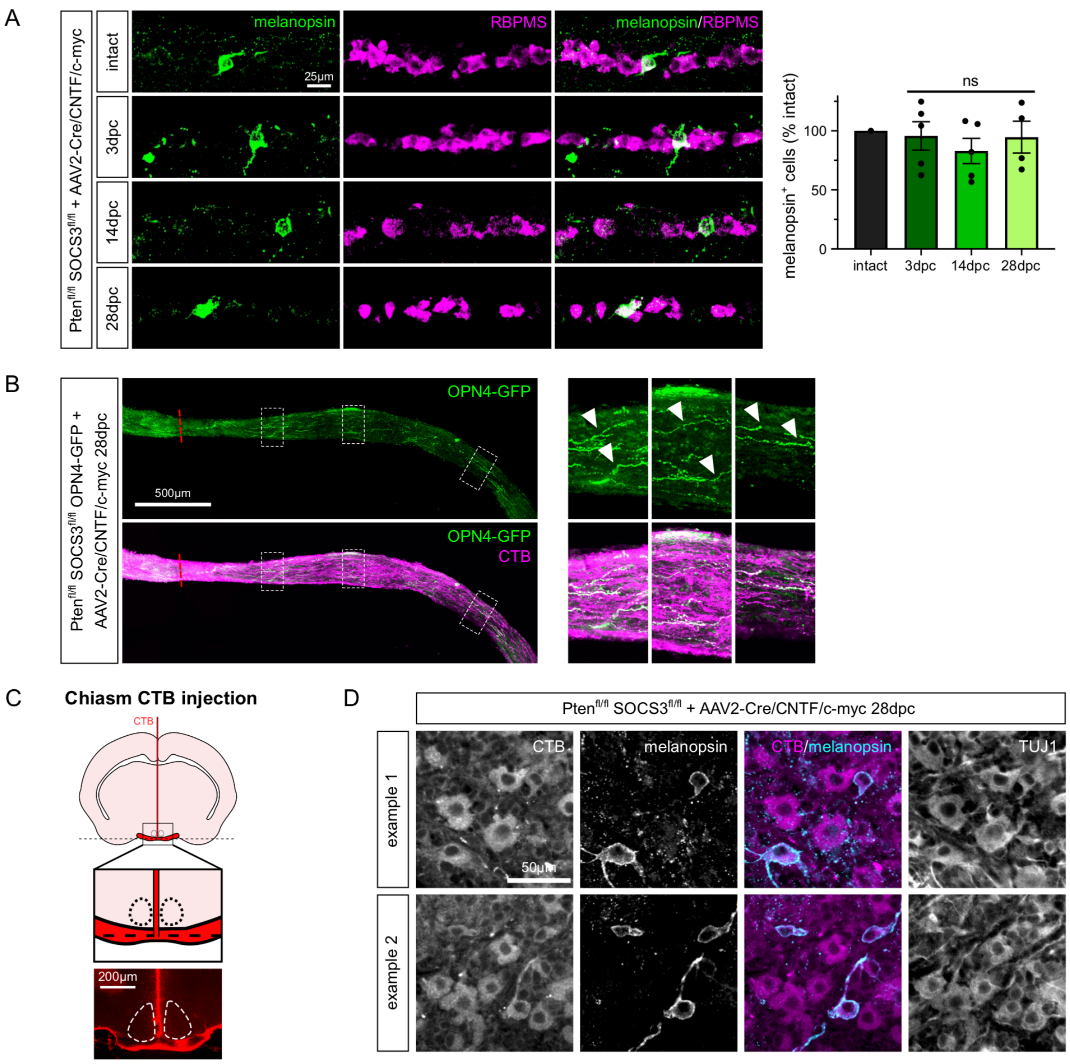


**Extended Data Figure 2: ipRGC survive and regenerate in the long-distance regeneration model.** (A) Representative confocal pictures of retina sections from Pten$^{fl/fl}$ SOCS3$^{fl/fl}$ eyes injected with AAV2-Cre/CNTF/c-myc in intact condition and at 3, 14 and 28dpc. ipRGC are labeled with anti-melanopsin antibody (green) and RGC are labeled with anti-RBPMS antibody (magenta). Corresponding quantification of the number of melanopsin$^{+}$ cells, in percentage of total ipRGC in intact condition. Data are expressed as mean +/- s.e.m. Kruskal-Wallis test, ns: not significant. (B) Representative confocal picture of optic nerve section from a Pten$^{fl/fl}$ SOCS3$^{fl/fl}$ OPN4-GFP mouse injected with AAV2-Cre/CNTF/c-myc at 28dpc. ipRGC axons are GFP$^{+}$ (green). Regenerating axons are traced with CTB (magenta). The red dashed line indicates the lesion site. Arrowheads point to GFP$^{+}$ regenerating axons distal to the lesion site. (C) Principle of stereotaxic injection of CTB in the optic chiasm for backtracing of regenerating axons. (D) Representative confocal pictures of whole-mount retinas labeled with anti-CTB antibody (magenta) and anti-melanopsin antibody (cyan).

### Extended Data Figure 3

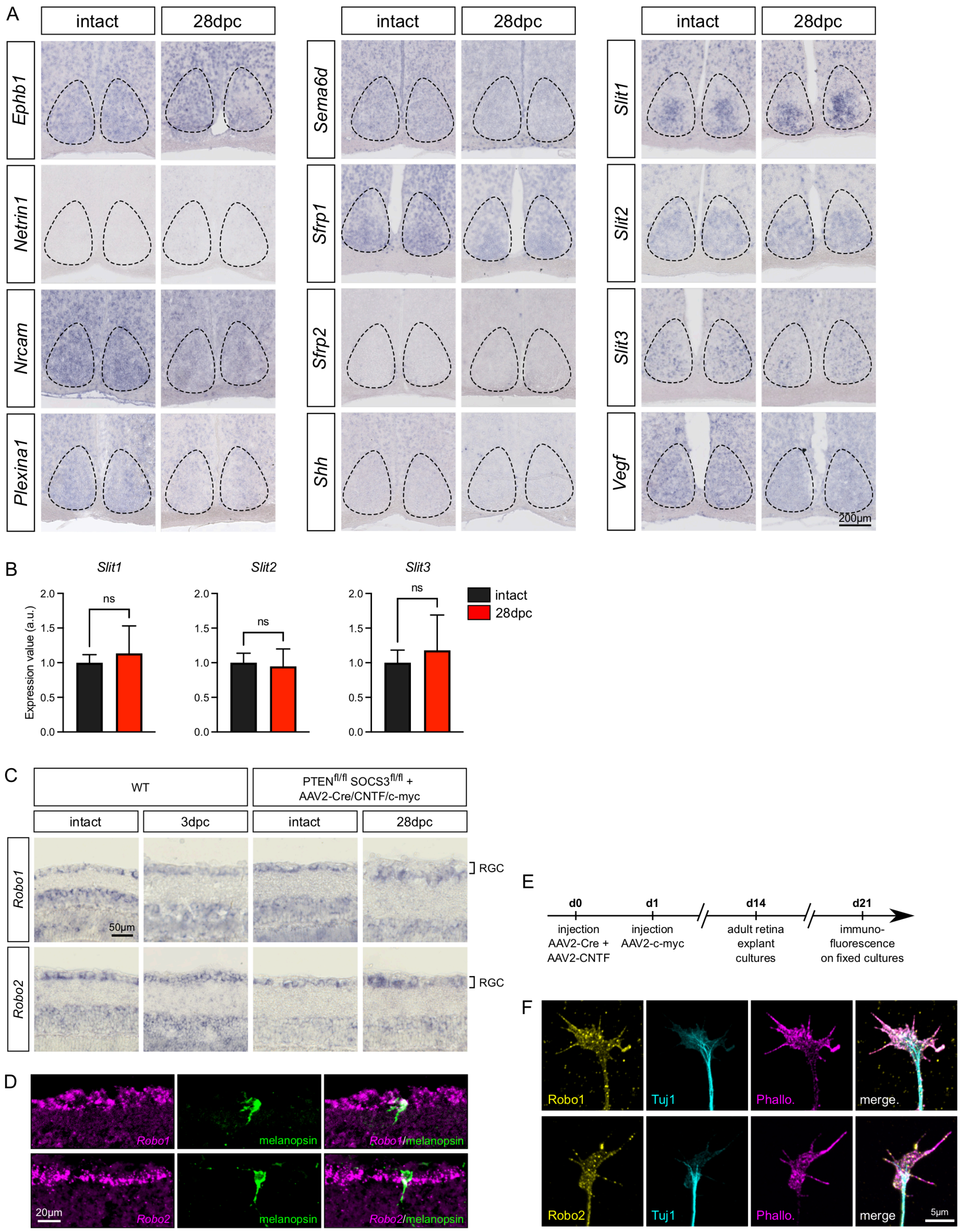


**Extended Data Figure 3: The SCN expresses Slit and the RGC express their corresponding receptors Robo.** (A) In situ hybridization showing guidance cues expression in the SCN of WT mice, in intact condition and at 28dpc. (B) Quantitative PCR on reverse transcription from WT SCN RNA, in intact condition and at 28dpc. Expression levels of *Slit1*, *Slit2* and *Slit3* normalized to *Gapdh*, relative to intact. Data are expressed as mean +/- s.e.m.

N=3-5 mice in each condition. Unpaired t-test, ns: not significant. (C) In situ hybridization showing *Robo1* and *Robo2* expression in RGC of WT mice, in intact condition and at 3dpc; and in RGC of Pten$^{fl/fl}$ SOCS3$^{fl/fl}$ mice injected with AAV2-Cre/CNTF/c-myc, in intact condition, at 3dpc and at 28dpc. (D) Representative confocal pictures showing *Robo1* and *Robo2* expression (in situ hybridization with TSA amplification, magenta) in ipRGC (labeled with anti-melanopsin antibody, green). (E) Timeline of adult retina explant cultures. (F) Representative confocal pictures with Airyscan technology of individual growth cones in an explant culture of a Pten$^{fl/fl}$ SOCS3$^{fl/fl}$ retina injected with AAV2-Cre/CNTF/c-myc. Robo1 and 2 (green) are expressed on growth cones, labeled with anti-β-tubulin III (TUJ1, cyan) and phalloidin (magenta).

### Extended Data Figure 4

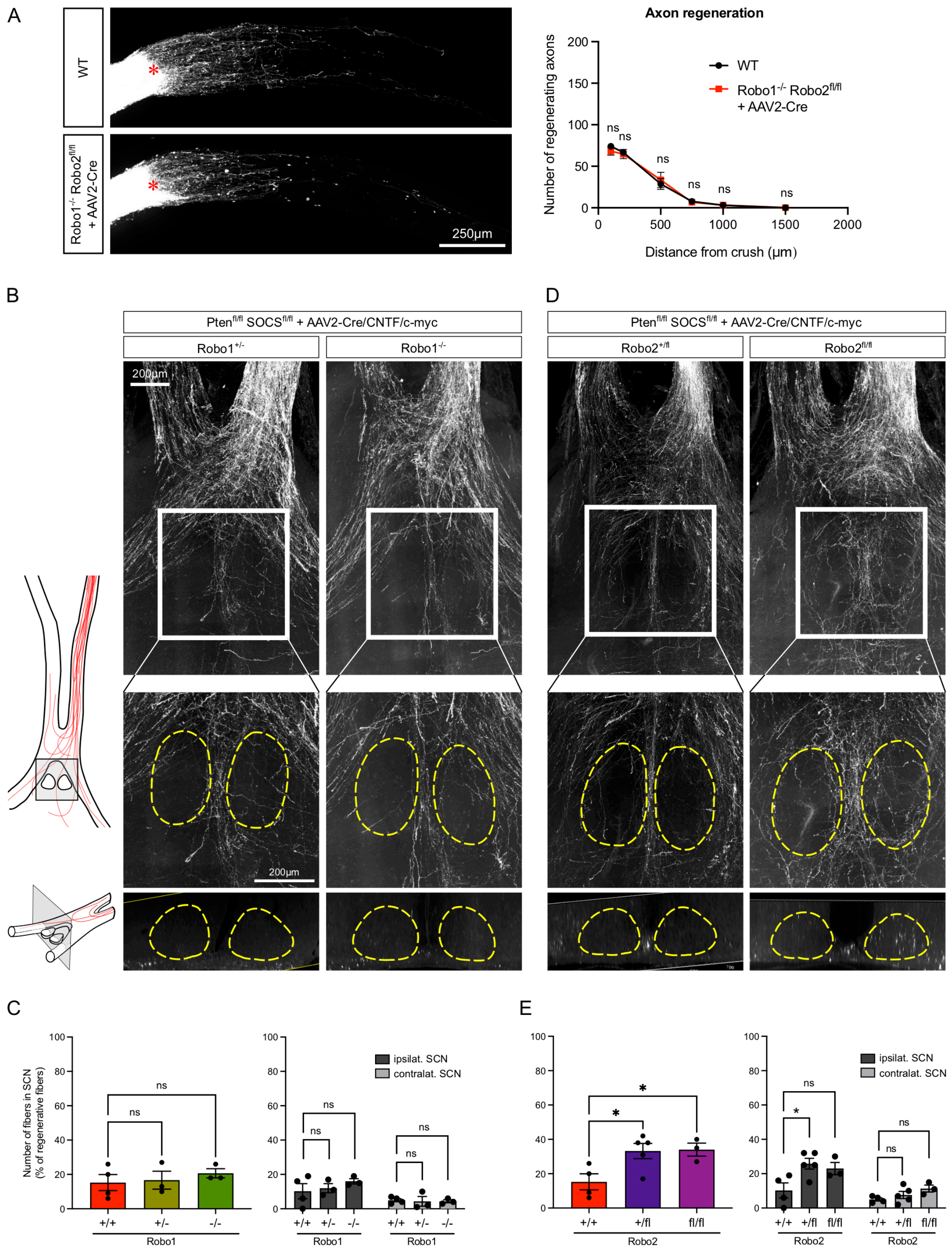


**Extended Data Figure 4: Regenerating axons in the optic chiasm and SCN region in single Robo-knockout conditions.** (A) Confocal pictures of whole optic nerves in WT and Robo1$^{-/-}$ Robo2$^{fl/fl}$ + AAV2-Cre conditions. Quantification of the number of regenerating axons at defined distances from the injury site (indicated with a red star). (B) Confocal picture of whole optic nerves and optic chiasm at 28dpc in Pten$^{fl/fl}$ SOCS3$^{fl/fl}$ Robo1$^{+/-}$ and Pten$^{fl/fl}$

SOCS3$^{fl/fl}$ Robo1$^{-/-}$. Regenerating axons are traced using CTB (white). Pictures are representive of N=3-5 animals in each condition. Zoom pictures: the SCN is indicated with the yellow dashed line on the maximum projection panel and on the XZ orthogonal section panel. (C) Quantification of the number of axons entering the SCN and distribution of axons entering the ipsilateral and the contralateral SCN. Data are expressed as mean +/- s.e.m. One-way ANOVA with Dunnett's correction, * p-value < 0.05, ns: not significant. (D) Confocal picture of whole optic nerves and optic chiasm at 28dpc in Pten$^{fl/fl}$ SOCS3$^{fl/fl}$ Robo2$^{+/fl}$ and Pten$^{fl/fl}$ SOCS3$^{fl/fl}$ Robo2$^{fl/fl}$. Regenerating axons are traced using CTB (white). Pictures are representive of N=3-5 animals in each condition. Zoom pictures: the SCN is indicated with the yellow dashed line on the maximum projection panel and on the XZ orthogonal section panel. (E) Quantification of the number of axons entering the SCN and distribution of axons entering the ipsilateral and the contralateral SCN. Data are expressed as mean +/- s.e.m. One-way ANOVA with Dunnett's correction, * p-value < 0.05, ns: not significant.

### Extended Data Figure 5

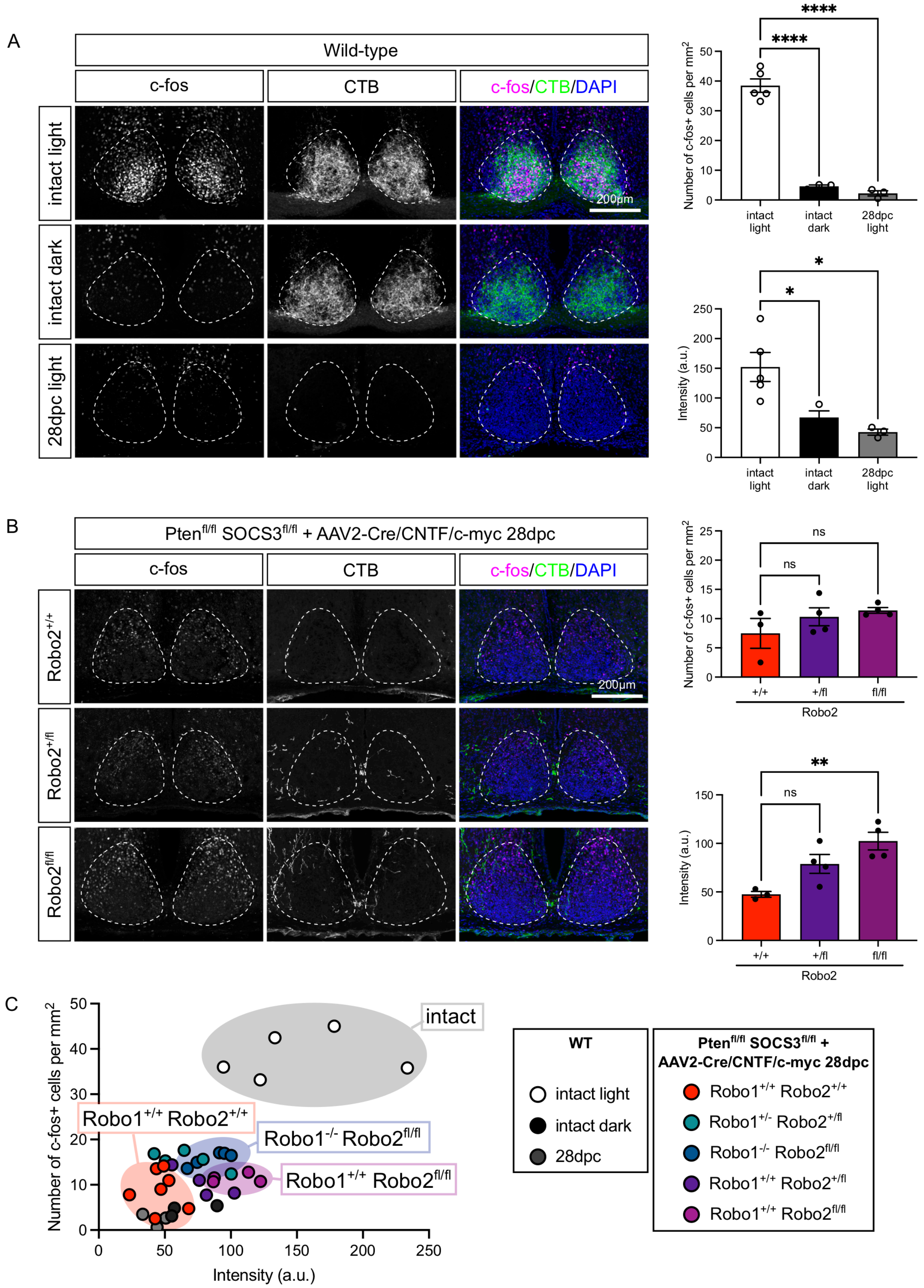


**Extended Data Figure 5: Neuronal activation assay in intact and regenerating conditions.** (A) Confocal pictures showing SCN expression of c-fos (magenta) in intact condition, where mice were subjected to light or left in the dark, and in control condition at

28dpc. The SCN is indicated with a white dashed line. Each picture is representative of N=3-5 mice. Corresponding quantification of c-fos$^{+}$ cell number per mm$^{2}$ section in the SCN (top) and of c-fos intensity in c-fos$^{+}$ cells in the SCN (bottom). Data are expressed as mean +/- s.e.m. One-way ANOVA with multiple comparisons, **** p-value < 0.0001. (B) Confocal pictures showing SCN expression of c-fos (magenta) in the 28dpc regeneration model, in Pten$^{fl/fl}$ SOCS3$^{fl/fl}$ Robo1$^{+/+}$ Robo2$^{+/+}$, Robo1$^{+/-}$ Robo2$^{+/fl}$ and Robo1$^{-/-}$ Robo2$^{fl/fl}$ conditions. CTB (green) was used to trace regenerating axons. The SCN is indicated with a white dashed line. Each picture is representative of N=3-4 mice. Corresponding quantification of c-fos$^{+}$ cell number per mm$^{2}$ section in the SCN (top) and of c-fos intensity in c-fos$^{+}$ cells in the SCN (bottom). Data are expressed as mean +/- s.e.m. One-way ANOVA with multiple comparisons, * p-value < 0.05. (C) Representation of c-fos$^{+}$ cell number per mm$^{2}$ section in the SCN versus c-fos intensity.

### Extended Data Figure 6

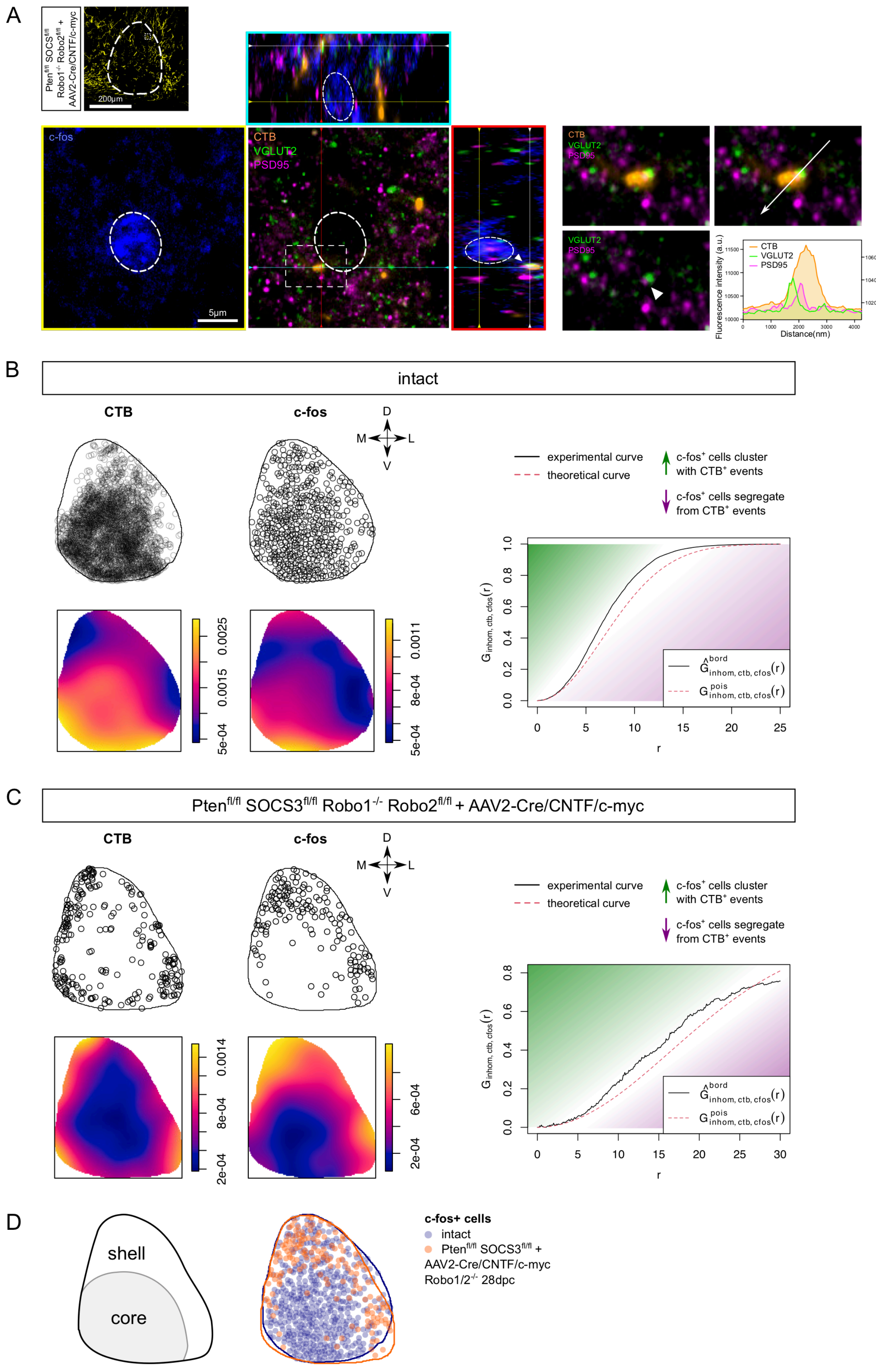

**Extended Data Figure 6: Neuronal activation in the SCN correlates with the distribution of regenerating axons entering the SNC.** (A) Left: confocal picture of the ipsilateral SCN from a Pten$^{fl/fl}$ SOCS3$^{fl/fl}$ Robo1$^{-/-}$ Robo2$^{fl/fl}$ mouse injected with AAV2-Cre/CNTF/c-myc with bilateral optic nerve crush at 28dpc. Centre: 3D confocal picture with Airyscan technology of an individual regenerative fiber forming a synapse in the SCN, adjacent to a c-fos$^{+}$ cell (blue). The pre-synaptic compartment marker VGLUT2 (green) colocalizes with the CTB$^{+}$ fiber (orange) and is adjacent to the post-synaptic compartment marker PSD95 (magenta). Right: zoom showing the synapse formed by the CTB$^{+}$ regenerative fiber. Profile of intensity in a single z plan along the white arrow. (B) Spatial point pattern analysis in an intact condition. Density maps are represented with pseudo-color. The right graph shows the G-cross function, i.e. the cumulative distribution of nearest neighbour distance from CTB$^{+}$ events to c-fos$^{+}$ cells. (C) Spatial point pattern analysis in Pten$^{fl/fl}$ SOCS3$^{fl/fl}$ Robo1$^{-/-}$ Robo2$^{fl/fl}$ mouse injected with AAV2-Cre/CNTF/c-myc at 28dpc. (D) Overlay of c-fos$^{+}$ events in intact versus regenerating condition. CTB$^{+}$ events and c-fos$^{+}$ cells cluster in the core of the SCN in intact condition, while CTB$^{+}$ events and c-fos$^{+}$ cells cluster in the shell of the SCN in the regenerating condition.

**Extended Data Figure 7**

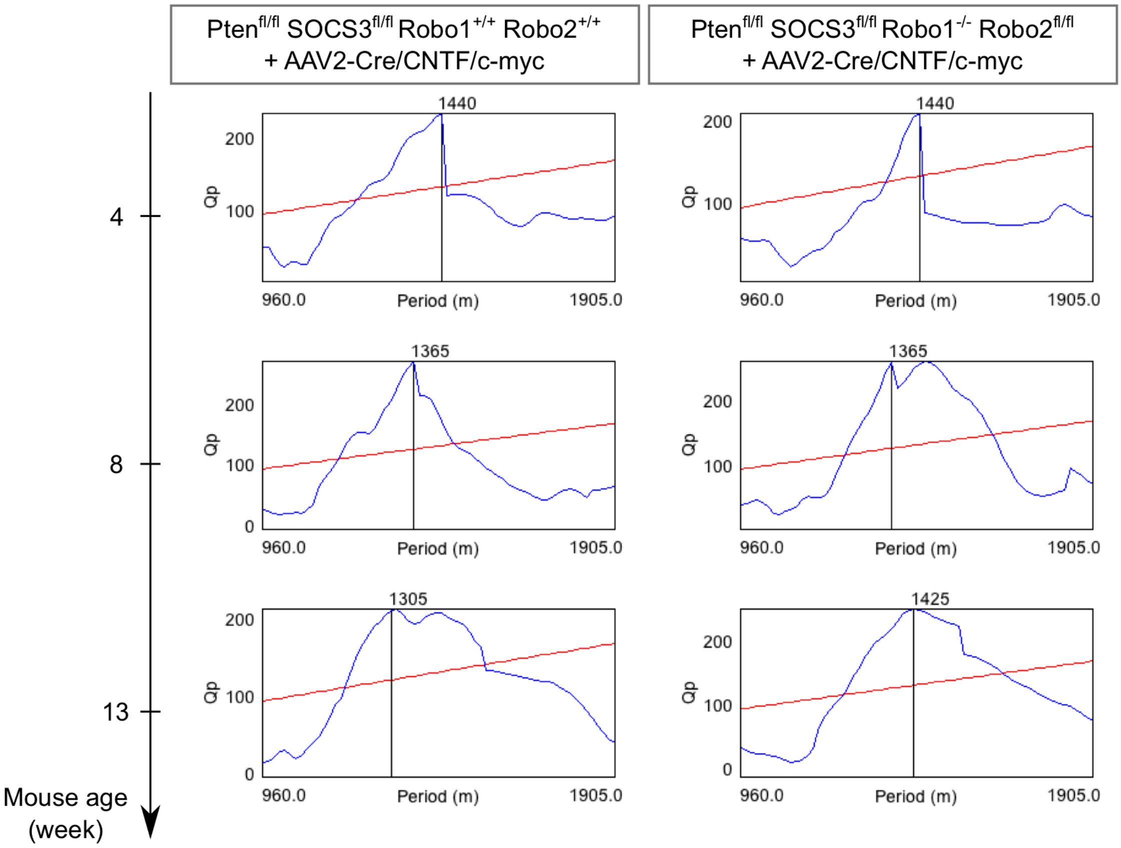


**Extended Data Figure 7: Mature axon guidance during long-distance regeneration leads to functional recovery of SCN activity.** Examples of periodogram showing Chi-2 periods (in min) in the control group (Pten$^{fl/fl}$ SOCS3$^{fl/fl}$ + AAV2-Cre/CNTF/c-myc) versus reinnervated SCN group (Pten$^{fl/fl}$ SOCS3$^{fl/fl}$ Robo1$^{-/-}$ Robo2$^{fl/fl}$ + AAV2-Cre/CNTF/c-myc). ONC is performed in 6-week-old mice, so mouse age week 4 is basal level, mouse age week 8 is 14dpc and mouse age week 13 is after 42dpc.

### Supplementary Movie captions

**Movie S1: Regenerating axons avoid entering the SCN in the long-distance regeneration model, despite sufficient growth in the SCN vicinity, related to Figure 4.** 3D imaging and reconstruction of $CTB^{+}$ regenerating axons in the SCN region at 28dpc, in $Pten^{fl/fl}$ $SOCS3^{fl/fl}$ $Robo1^{+/+}$ $Robo2^{+/+}$ + AAV2-Cre/CNTF/c-myc condition.

**Movie S2: Modulation of Robo1 and Robo2 make regenerating axons enter and stay in the SCN, related to Figure 4.** 3D imaging and reconstruction of $CTB^{+}$ regenerating axons in the SCN region at 28dpc, in $Pten^{fl/fl}$ $SOCS3^{fl/fl}$ $Robo1^{-/-}$ $Robo2^{fl/fl}$ + AAV2-Cre/CNTF/c-myc condition.

### Resource table for Material and Methods

| REAGENT or RESOURCE | SOURCE | IDENTIFIER |
|---|---|---|
| Antibodies | | |
| Chicken polyclonal anti-GFP antibody | Abcam | Cat# ab13970; RRID: AB_300798 |
| Goat polyclonal anti-VGLUT2 antibody | Abcam | Cat # ab178538 |
| Guinea polyclonal pig anti-RBPMS antibody | Millipore | Cat# ABN1376; RRID: AB_2687403 |
| Mouse monoclonal anti-tubulin β3 (TUJ1) antibody | Biolegend | Cat# 801202; RRID: AB_10063408 |
| Mouse monoclonal anti-PSD95 antibody | Abcam | Cat# ab2723 ; RRID : AB_303248 |
| Rabbit polyclonal anti-VIP (Vasoactive Intestinal Peptide) antibody | Immunostar | Cat# 20077; RRID: AB_572270 |
| Rabbit monoclonal anti-c-fos antibody | Cell Signaling Technology | Cat# 2250 ; RRID : AB_2247211 |
| Rabbit polyclonal anti-melanopsin antibody | Abcam | Cat# ab19306; RRID: AB_444842 |
| Goat polyclonal anti-Robo1 antibody | R and D Systems | Cat# AF1749; RRID: AB_354969 |

| | | |
|---|---|---|
| Goat polyclonal anti-Robo2 antibody | Dr Alain Chédotal; Dominici et al., 2018 | |
| Donkey anti-rabbit IgG (H+L) highly cross-absorbed secondary antibody, Alexa Fluor 488 | Thermo Fisher Scientific | Cat# A-21206; RRID: AB_2535792 |
| Donkey anti-rabbit IgG (H+L) highly cross-absorbed secondary antibody, Alexa Fluor 647 | Thermo Fisher Scientific | Cat# A-31573; RRID: AB_2536183 |
| Donkey anti-chicken IgY (IgG) (H+L) antibody, Alexa Fluor 488 | Jackson ImmunoResearch Labs | Cat# 703-545-155; RRID: AB_2340375 |
| Donkey anti-guinea pig IgG (H+L), Alexa Fluor 647 | Jackson ImmunoResearch Labs | Cat# 706-605-148; RRID: AB_2340476 |
| Donkey anti-goat IgG (H+L), Alexa Fluor 488 | Thermo Fisher Scientific | Cat# A11055 ; RRID : AB_2534102 |
| Donkey anti-mouse IgG (H+L), Alexa Fluor 647 | Thermo Fisher Scientific | Cat# A31571 ; RRID_162542 |
| Donkey anti-mouse IgG (H+L), Alexa Fluor 568 | Thermo Fisher Scientific | Cat# A10037 ; RRID : AB_2534013 |
| Bacterial and virus strains | | |
| AAV2-Cre | Belin et al., 2015 | N/A |
| AAV2-CNTF | Belin et al., 2015 | N/A |
| AAV2-c-myc | Belin et al., 2015 | N/A |
| Chemicals, peptides, and recombinant proteins | | |
| Cholera toxin B subunit (CTB)-Alexa 488 | Thermo Fisher Scientific | Cat# C34775 |
| Cholera toxin B subunit (CTB)-Alexa 555 | Thermo Fisher Scientific | Cat# C22843 |
| Cholera toxin B subunit (CTB)-Alexa 647 | Thermo Fisher Scientific | Cat# C34778 |
| Laminin | Sigma Aldrich | Cat# L2020 |
| Poly-L-Lysine | Sigma Aldrich | Cat# P1399 |
| Collagen I, rat tail | Thermo Fisher Scientific | Cat# A1048301 |
| Experimental models: Organisms/strains | | |
| Mouse: Pten$^{fl/fl}$ SOCS3$^{fl/fl}$ | Sun et al., 2011; The Jackson Laboratory | N/A |

| | | |
|---|---|---|
| Mouse: OPN4-GFP $Pten^{fl/fl}$ $SOCS3^{fl/fl}$ | Schmidt et al., 2008; Sun et al., 2011; The Jackson Laboratory | N/A |
| Mouse: $Pten^{fl/fl}$ $SOCS3^{fl/fl}$ $Robo1^{-/-}$ $Robo2^{fl/fl}$ | Long et al., 2004; Lu et al., 2007; Sun et al., 2011; Dr Alain Chédotal | N/A |
| Mouse: $Pten^{fl/fl}$ $SOCS3^{fl/fl}$ $Robo1^{-/-}$ | Long et al., 2004; Lu et al., 2007; Sun et al., 2011; Dr Alain Chédotal | N/A |
| Mouse: $Pten^{fl/fl}$ $SOCS3^{fl/fl}$ $Robo2^{fl/fl}$ | Long et al., 2004; Lu et al., 2007; Sun et al., 2011; Dr Alain Chédotal | N/A |
| Mouse: OPN4-GFP $Pten^{fl/fl}$ $SOCS3^{fl/fl}$ $Robo1^{-/-}$ $Robo2^{fl/fl}$ | Schmidt et al., 2008; Sun et al., 2011; The Jackson Laboratory; Long et al., 2004; Lu et al., 2007; Sun et al., 2011; Dr Alain Chédotal | N/A |
| Mouse: $Slit1^{-/-}$ | Dr Alain Chédotal | N/A |
| Mouse: $Slit2^{fl/fl}$ | Dr Alain Chédotal | N/A |
| Mouse: $Slit1^{-/-}$ $Slit2^{fl/fl}$ | Dr Alain Chédotal | N/A |
| Mouse: $Slit3^{-/-}$ | Dr Alain Chédotal | N/A |
| Mouse: ROSA-TdTomato floxed | The Jackson Laboratory | RRID:IMSR_JAX:007914 |
| Oligonucleotides | | |
| Primer ISH: Plexin-A1 Forward 5’-CCCCCACTGTGGCTGGTGTG-3’ | This paper | N/A |
| Primer ISH: Plexin-A1 Reverse 5’-GAAAGGCGCAGTCAGCCGCA-3’ | This paper | N/A |
| Primer ISH: Robo1 Forward 5’-AGCAGTGGATGGCACTTTAA-3’ | This paper | N/A |

| | | |
|---|---|---|
| Primer ISH: Robo1 Reverse<br>5’-GGAAAAGGTAAATGGCGTTA-3’ | This paper | N/A |
| Primer ISH: Robo2 Forward<br>5’-CTTTTCCCGAATCAACCTCA-3’ | This paper | N/A |
| Primer ISH: Robo2 Reverse<br>5’-GGGAGGTCATTCATATCATA-3’ | This paper | N/A |
| Primer ISH: Sema6D Forward<br>5’-TTCCTCCATGTGTGTCCTGT-3’ | This paper | N/A |
| Primer ISH: Sema6D Reverse<br>5’-GTTGGGTAAATGACTGGGTGATGT-3’ | This paper | N/A |
| Primer ISH: Sfrp1 Forward<br>5’-GGACCTGAGGCTGTGCCACA-3’ | This paper | N/A |
| Primer ISH: Sfrp1 Reverse<br>5’-TCTTCTTGGGGACAATCTTC-3’ | This paper | N/A |
| Primer ISH: Sfrp2 Forward<br>5’-GCCCAACCTGCTGGGCCACG-3’ | This paper | N/A |
| Primer ISH: Sfrp2 Reverse<br>5’-CGCCGTTCAGCTTGTAAATG-3’ | This paper | N/A |
| Primer ISH: Slit1 Forward<br>5’-CACTGGGTTGTTTAAGAAGC-3’ | This paper | N/A |
| Primer ISH: Slit1 Reverse<br>5’-CCTGGCCTTCCTCACACCTG-3’ | This paper | N/A |
| Primer ISH: Slit2 Forward<br>5’-CCTGCCAGCATGACTCCAAG-3’ | This paper | N/A |
| Primer ISH: Slit2 Reverse<br>5’-TCTATAGAGTTCCACGGCAA-3’ | This paper | N/A |
| Primer ISH: VEGF-A Forward<br>5'-GCAGCGACAAGGCAGACTA-3’ | This paper | N/A |
| Primer ISH: VEGF-A Reverse<br>5’-GCTAGCACTTCTCCCAGCTC-3’ | This paper | N/A |
| Primer qPCR: Slit1 Forward<br>5’-AGGTGCAAAAGGGCGAAT-3’ | This paper | N/A |
| Primer qPCR: Slit1 Reverse<br>5’-CGAGAGGGTACAGGCAGGT-3’ | This paper | N/A |

| | | |
|---|---|---|
| Primer qPCR: Slit2 Forward<br>5’-CGGGGACAGCTGTGATAGAG-3’ | This paper | N/A |
| Primer qPCR: Slit2 Reverse<br>5’-CCAAGCGAGATACTTTCTTAGTTGT-3’ | This paper | N/A |
| Primer qPCR: Slit3 Forward<br>5’-GCCACCTCAGTGAGAACCTC-3’ | This paper | N/A |
| Primer qPCR: Slit3 Reverse<br>5’-TGTCCCTCAAAGCCCAGA-3’ | This paper | N/A |
| Primer qPCR: Gapdh Forward<br>5’-GCATGGCCTTCCGTGTTC-3’, | This paper | N/A |
| Primer qPCR: Gapdh Reverse<br>5’-TGTCATCATACTTGGCAGGTTTCT-3’ | This paper | N/A |
| Recombinant DNA | | |
| Plasmid ISH: Ephrin-B1 | Dr Valérie Castellani | N/A |
| Plasmid ISH: Netrin1 | Dr Valérie Castellani;<br>Nawabi et al., 2010 | N/A |
| Plasmid ISH: NrCAM | Dr Valérie Castellani;<br>Nawabi et al., 2010 | N/A |
| Plasmid ISH: Shh | Dr Valérie Castellani<br>Charoy et al., 2012 | N/A |
| Plasmid ISH: Slit3 | Dr Alain Chédotal | N/A |
| Software and algorithms | | |
| Zen | Zeiss | N/A |
| Fiji | Image J | N/A |
| IMARIS v.9.6 | Bitplane | N/A |
| GraphPad Prism 9 | GraphPad software | N/A |
| Fusion | Andor | N/A |
